# Force-regulated chaperone activity of BiP/ERdj3 is opposite to their homologs DnaK/DnaJ: explained by strain energy

**DOI:** 10.1101/2023.03.16.532907

**Authors:** Souradeep Banerjee, Debojyoti Chowdhury, Soham Chakraborty, Shubhasis Haldar

## Abstract

Polypeptide chains experiences mechanical tension while translocating through cellular tunnel. In this scenario, interaction of tunnel-associated chaperones with the emerging polypeptide occurs under force; however, this force-regulated chaperone behaviour is not fully understood.

We studied the mechanical chaperone activity of two tunnel-associated chaperones BiP and ERdj3 both in the absence and presence of force; and compared to their respective cytoplasmic homologs DnaK and DnaJ. We found that BiP/ERdj3 shows strong foldase activity under force; whereas their cytoplasmic homolog DnaK/DnaJ behave as holdase. Importantly, these tunnel-associated chaperones (BiP/ERdj3) revert to holdase in the absence of force, suggesting that mechanical chaperone activity differs depending on the presence or absence of force. This tunnel-associated chaperone-driven folding event generates additional mechanical energy of up to 54 zJ that could help protein translocation. The mechanical-chaperone behaviour can be explained by strain theory: chaperones with higher intrinsic deformability function as mechanical foldase (BiP, ERdj3), while chaperones with lower intrinsic deformability act as holdase (DnaK and DnaJ). Our study thus unveils the underlying mechanism of mechanically regulated chaperoning activity and provides a novel mechanism of co-translocational protein folding.

**Significance:** The mechanical-activity of chaperones, located at the edge of a tunnel, could be different from their cytoplasmic homologs. Translocating substrates within the tunnel are known to experience mechanical constraints, whereas the cytosolic substrates interact with the chaperones in the absence of force.

To understand this phenomenon, we investigated two tunnel-associated chaperones BiP/ERdj3 and their cytosolic homologs-DnaK/DnaJ. We observed that BiP/ERdj3 possess strong foldase activity while their substrates are under force; whereas DnaK/DnaJ possess holdase function. Notably all these chaperones function as holdase in the absence of force, which suggest that mechanical chaperone activity is different with and without force. We explained this mechanical behaviour using strain theory, providing a physical mechanism of chaperone-assisted co-translocational protein folding.

## Introduction

Cotranslational protein folding events like protein translocation across the membrane or folding at the edge of a tunnel, produces pulling forces on the emerging polypeptide chain (1–3). Studies have also revealed that when nascent chains are translocated through SecYEG or ribosomal-exit tunnel near the membrane, they are under mechanical tension due to the confined geometry of the tunnel and electrostatic force among the residues (4, 5). Molecular chaperones, known to assist the protein folding through different ways, are located at the edge of those tunnels to interact with the nascent chain and influence the force transmission through them (6, 7). Notably, the tunnel-associated chaperone activity under force might be different than the cytosolic chaperones; as in the cytosol, the chaperones moves freely and interact with their substrate without any mechanical constraints. However, determination of the underlying source for this generic mechanical activity is limited in the literature.

To address this mechanical chaperone behaviour, we have selected two cytosolic chaperones DnaK and DnaJ; and their tunnel-associated homologs Binding immunoglobulin Protein (BiP) and Endoplasmic Reticulum DnaJ homologue 3 (ERdj3) to monitor their mechanical effect on protein folding dynamics using magnetic tweezers (8). The force-clamp methodology in magnetic tweezers allows us to specifically apply a constant force on the substrate without perturbing the interacting chaperones, for probing their activity under equilibrium conditions (9). More importantly, we can introduce different chaperones, either separately or in combination, into the experimental buffer to track their individual or collective effect on the folding dynamics of a single client protein molecule.

We have systematically investigated the mechanical activity of different chaperones on protein L, which has been previously used as a model chaperone-substrate in many force-spectroscopic studies (10–13). Protein L do not form any misfolded or intermediate state in the folding pathway and thus, possess two-state folding pathway (14, 15). Here we showed that cytosolic holdase chaperones behave differently while located at the edge of different molecular tunnels, including ER Sec61. Prototypical chaperones DnaK and DnaJ act as mechanical holdase; whereas their ER tunnel-associated variants BiP and ERdj3 possess strong foldase activity under force, producing a work-output of upto ∼54 zJ which could aid in the pulling of polypeptide from the confined tunnel and prevents backsliding of the polypeptide. By contrast, at near-zero force, these chaperones possess holdase activity similar to DnaK/DnaJ, plausibly indicating that mechanical activity of these chaperones is uniquely different from their foldase functions. Additionally, we explained the underlying source of this mechanical chaperone behaviour using strain theory, suggesting that the mechanical behaviour of chaperones arises from their intrinsic strain energy: DnaK and DnaJ have higher strain energy difference during their interaction with the unfolded protein L and thereby stabilizing the unfolded state; while BiP and ERdj3 exhibit higher energy difference during binding to folded protein L conformation, signifying favored folded state of protein L in their presence. Overall, these data suggest that chaperones, by modifying the mechanical folding landscape of substrate protein, modulate the mechanical energy; providing a distinct mechanistic view of mechanical chaperone behaviour during protein folding under force.

## Results

### Measurement of protein L folding dynamics by magnetic tweezers

We have performed the magnetic-tweezers experiment on a protein L octamer construct containing its B1 domains in tandem repeats, bracketed within N-terminal HaloTag and C-terminal AviTag, which are used for construct attachment to glass surface and paramagnetic bead, respectively (9, 10, 16). Applied force on paramagnetic beads can be determined using a calibration curve as an inverse function of the distance between permanent magnets and glass surface such that decreasing the distance would increase the applied force (Fig. 1b) (9). Detailed information about the force calibration method has been described previously (9, 10, 13, 16–19). Fig. 1a shows a typical magnetic tweezers trajectory: first the polyprotein is unfolded at 45 pN force, resulting in eight distinct stepwise extension of ∼15 nm. Next, we quenched the pulse to 6 pN, where the polyprotein first collapses and then completely refolds; followed by a balanced folding-unfolding transition at an equilibrium condition. Minimum time taken for these complete unfolding and refolding is defined as first passage time (FPT) of unfolding and refolding, respectively. Averaging such FPT values of numerous trajectories are defined as Mean-FPT (MFPT), which has been previously used as model-free metric for protein folding kinetics (20–22) (Eq.1).

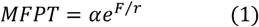

Where, *α* is a time fitting parameter and *r* is a force fitting parameter, as described previously (10, 23). For MFPT unfolding, r has a negative value as MFPT of unfolding declines with the force. Next, we determined the folding probability (FP) of protein L from the equilibrium phase of several trajectories using dwell-time analysis within 2-12 pN force and fitted to the sigmoid equation (Eq. 2) (10, 13, 17, 18, 24):

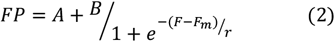

Where *A* and *B* are base and maximum value of the plot, *F*_*m*_ is half-point force and *r* is rate.

**Figure 1:**
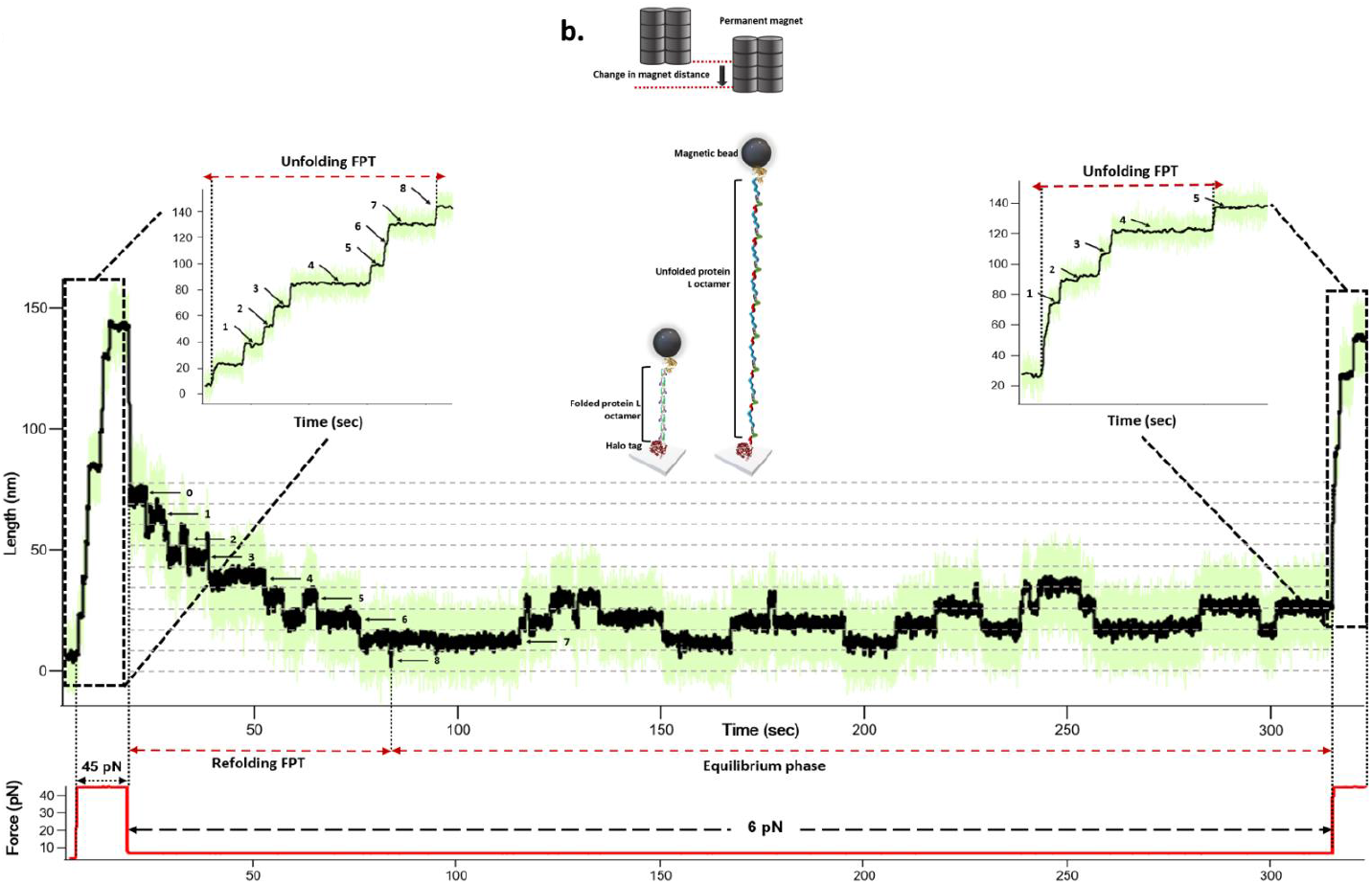
Representative trajectory protein L octamer obtained from magnetic tweezers: **(a) Representative trajectory of protein L:** Using the force-clamp technology the polyprotein construct is initially unfolded at a high force pulse of 45 pN, which is observed as eight distinct unfolding steps after the initial extension (inset); followed by a quench pulse at 6 pN. During quenching or refolding phase, the polyprotein first experiences an elastic contraction, followed by sequential refolding of all the domains and at last attains the dynamic equilibrium phase. The minimum time required for the complete unfolding of the polyprotein is the unfolding FPT (left inset); and similarly, the minimum time taken for complete refolding is called refolding FPT. To crosscheck proper refolding of the domains, a probe pulse has been applied after refolding phase to extend the protein and observed that the number of unfolded steps is matching exactly with the number of domains refolded. The FP is calculated from the equilibrium dynamics displayed by the protein construct. (**b) Experimental setup:** An octamer construct of protein L domain is biotinylated at its C-terminal AviTag and attached to the paramagnetic bead using biotin-streptavidin links; while the construct is attached to the glass surface through N-terminal HaloTag chemistry. Applied force is controlled by a pair of permanent magnets and is an inverse function of the distance between the paramagnetic bead and permanent magnet. Application of higher force results in the sequential unfolding of all eight domains in the construct (Figure is not scaled).

### Comparative effect of chaperones on protein L folding dynamics

Fig. 2 shows variation in the protein L FP against the force both in the absence and presence of BiP and ERdj3. We tested the chaperone activity of BiP and ERdj3 at a concentration, where their mechanical effect become saturated; and for DnaK and DnaJ, the same concentration has been used for the comparison with their homologs (Supplementary Fig. 1 and 2). Previously, it has been shown that chaperones, both with and without the force, can potentially interact with the partially-folded or near-native, folded and collapsed state of substrates(25–32); however, to further investigate its effects on protein folding under mechanical constraint, we have measured unfolding and refolding kinetics, and folding probability of protein L. Interestingly, we found that both BiP complex (BiP+ERdj3+Sil1+ATP), and its individual chaperone components BiP and ERdj3 shift the protein L half-point force (defined as a force where FP is equal to 0.5) towards higher force (Fig. 2 and Supplementary Fig. 3 and 4). By contrast, their cytoplasmic homolog DnaK/DnaJ individually moves the FP towards the lower force range with no changes while working synergistically (Fig. 2; Supplementary Fig. 5). Additionally, we plotted the FP curve difference with and without chaperones; showing that the mechanical effect of these chaperones is most prominent at ∼7 pN (Supplementary Fig. 6). To characterize the folding mechanism in detail, we measured both unfolding kinetics and refolding kinetics with BiP and ERdj3; and observed that these tunnel-associated homologs decreases the unfolding and increases the refolding kinetics of protein L under force (Supplementary Fig. 7-12), suggesting their interaction with both the folded and unfolded protein L states. Contrastingly, their cytoplasmic homologs inversely modulate the folding kinetics of protein L (Supplementary Fig. 13-16). The folding probability and kinetics data combinedly indicate that BiP and its cochaperone ERdj3 reshape the mechanical folding landscape of protein L towards the folded state through destabilizing the unfolded state and stabilizing the refolded state of protein L.

**Figure 2:**
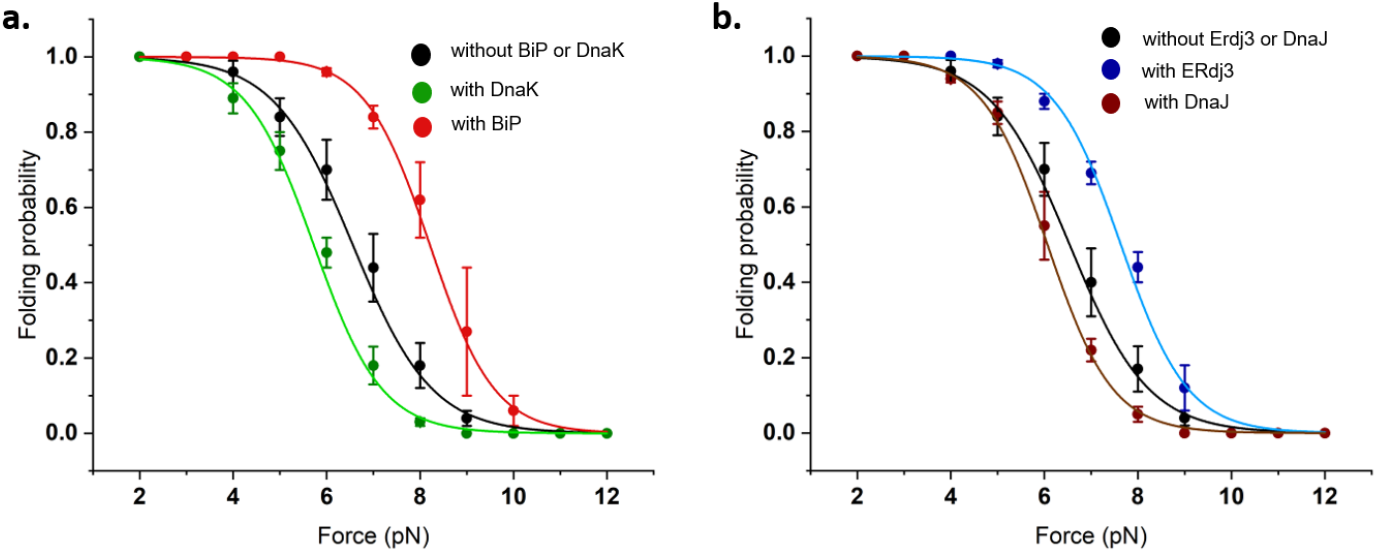
Comparative effect of tunnel-associated chaperones and their cytosolic holdases on protein L folding probability: **(a)Effect of BiP and DnaK:** Folding probability (FP) of protein L is plotted against the force. In the absence of any chaperones (control, black circles), protein L exhibits a half-point force of 6.4 pN, which has been observed to upshift to 8.2 pN in the presence of BiP (red circles). However, DnaK change the FP towards ∼5.9 pN. **(b) Folding probability in presence of ERdj3 and DnaJ:** ERdj3 positively modulates the folding probability of protein L by increasing half-point force to 7.7 pN; whereas its cytosolic homolog DnaK downshifts the half-point force to 5.7 pN. For each data points, n≥5 molecules were considered. Each FP data were observed for >250 s. Error bars represent standard error of mean (s.e.m).

### Effect of BiP and ERdj3 in near-zero force range

The mechanical foldase activity of BiP/ERdj3, while their substrate is under force, is independent of their well-established role of stabilizing the unfolded state; however, these experiments were mostly performed in the absence of force (8, 33, 34). To correlate their finding with our results, we further investigate the effect of these chaperones on protein L folding in near-zero force regime (0.3 pN), we designed a pulse protocol: firstly, the octamer construct was completely unfolded by a fingerprint pulse, followed by quenching it to near-zero range (0.3 pN) for different durations. Subsequently, a probe pulse was applied to monitor how many domains become folded during quench time (Fig. 4a). We reported the percentage of folded domain against the quench time with and without the BiP and ERdj3 (Fig. 4b and 4c). We found that both BiP and ERdj3 prevent the protein L refolding in near-zero force and thus, function as strong holdase chaperones. This result coincides with previous studies that BiP and ERdj3, very similar to their respective homologs DnaK and DnaJ, binds to and stabilizes the unfolded state of the substrate (35).

### Mechanical workdone by chaperones during protein L folding

Protein folding at the edge of any molecular tunnel generates mechanical work; and tunnel-associated chaperones could influence the folding ability through interacting with substrates under force(10, 19). This mechanical work during protein folding could be estimated by multiplying the force with step sizes, as described previously (18, 36). To check whether there is any intermediate or molten-globule like states of protein L, we also measured their step size while monitoring the folding events under force. However, since protein L do not form any intermediate or molten-globule like state, no altered step sizes have been observed during the folding events and their differences are non-significant in case of different chaperones (Supplementary Fig. 17). This implies protein folding at higher forces delivers more mechanical work done; however, at this regime, FP of a protein also decreases. Therefore, a more precise estimation of workdone is achieved by multiplying FP with folding work at a certain force. We found that the chaperones profoundly modulate the mechanical workdone within an equilibrium force range of 5-10 pN. Although chaperones do not change the step size (Supplementary Fig. 17), they significantly shift the FP curve and consequently, the mechanical workdone during protein L folding. While DnaK or DnaJ halves the workdone at 7 pN; BiP and ERdj3, individually or combinedly as a complex, increases the workdone by ∼1.5 to 2.5 folds, showing their prominent foldase activity under force (Fig. 3a and 3b; Supplementary Fig. 18).

**Figure 3:**
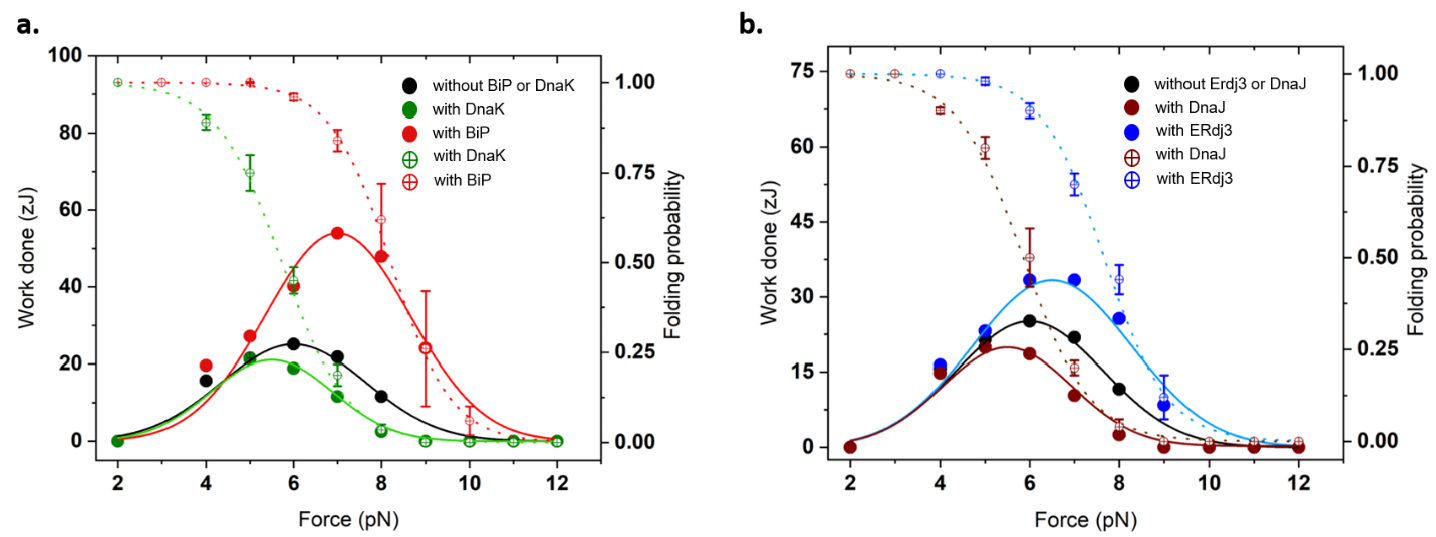
Chaperones modulate the mechanical work done of protein L folding: The mechanical workdone by the protein L folding is measured as a product of folding work with the folding probability (round circles with cross). The expected folding work is estimated by multiplying the step size of protein L with respective force. (a) **Work done in presence of BiP and DnaK:** In the absence of any chaperone (control, black circles), protein L folding delivers an optimal mechanical work done of 22±0.1 zJ at 7 pN; whereas, in the presence of 25 μM BiP, this work output is increased to 54±0.04 zJ (red circles). By contrast, DnaK decreases the mechanical work done to 11.6±0.04 zJ (green circles). **(b)Work done in presence of ERdj3 and DnaJ:** Similarly, at 7pN, DnaJ as a holdase chaperone decreases the mechanical workdone to 10.3±0.01 zJ; however, its counterpart ERdj3 delivers 33.3±0.02 zJ of mechanical energy during protein L folding. Error bars represent s.e.m.

**Figure 4:**
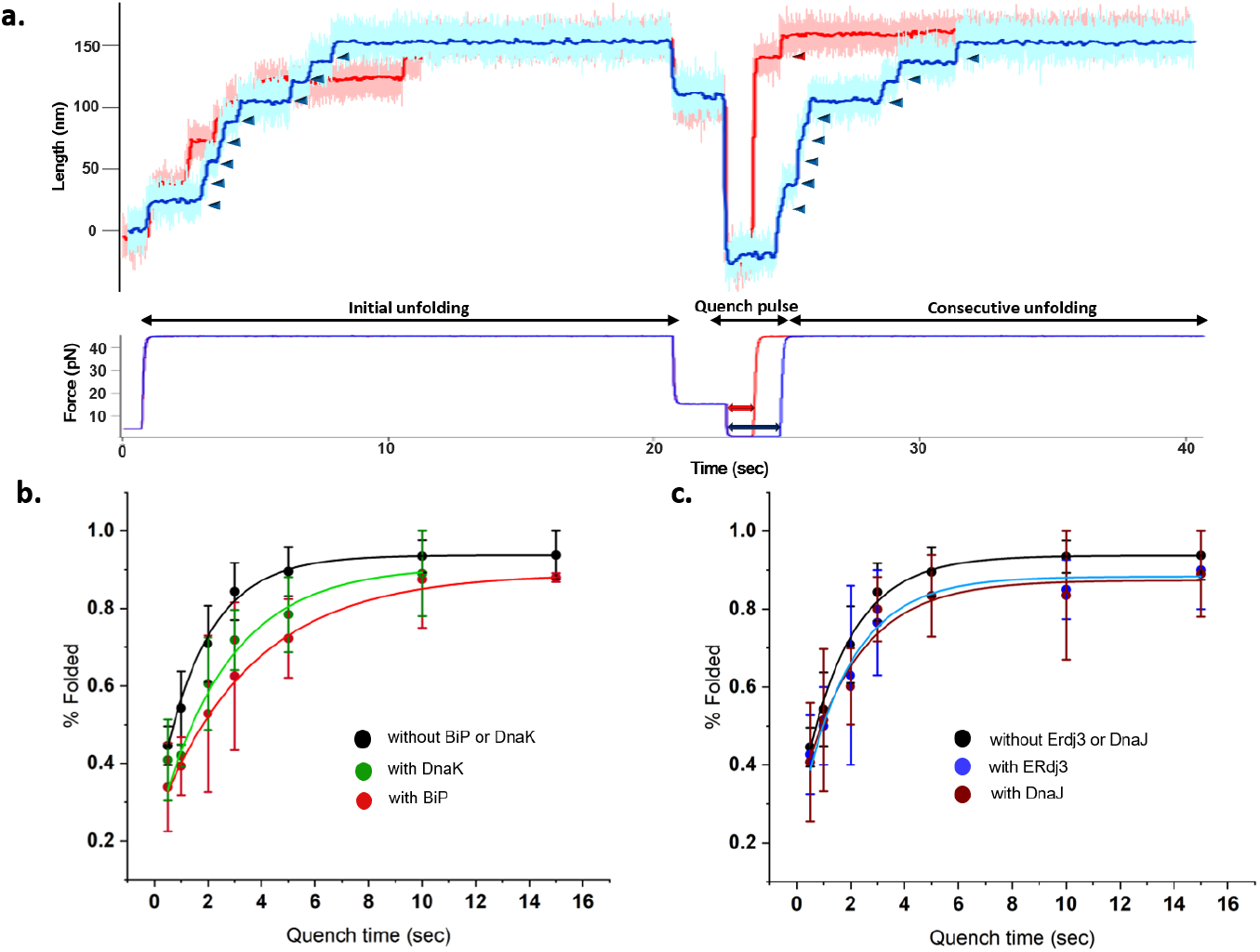
Activity of mechanical chaperones at near zero force regime after quench pulses: **(a) Protocol used for monitoring activity of mechanical chaperones at near zero force (0.3 pN):** First, the protein L octamer is unfolded at 45 pN for 20 s, followed by a quench pulse of 15 pN for 2 s. Thereafter, the polyprotein was maintained at 0.3 pN through different quench times (0.5, 1, 2, 3, 5, 10 and 15 s). Representative traces display eight distinct unfolding steps (blue arrows) at 45 pN, followed by two different quench times i.e. 1 s and 3 s. Again, a probe pulse of 45 pN shows only one unfolding step (red arrow) for 1 s quench time; whereas all eight domains unfold in a 3 s. **(b) Refolded percentage of protein L in presence of BiP and DnaK:** By fitting to a single exponential equation, the refolded percentage of native protein L has been found to be 0.6±0.1; while in presence of 25 μM BiP and DnaK, the folding rate is reduced to 0.3±0.04 and 0.4±0.1, respectively. Each datapoint represents n≥4 molecules. Error bars represent s.e.m. **(c) Refolded percentage of protein L with ERdj3 and DnaJ:** Percentage of refolded domains have also been observed to decrease with 10 μM ERdj3 (blue) and DnaJ (brown) in different quench time at 0.3 pN to 0.5±0.1 and 0.5±0.1 respectively. Each datapoint represents n≥4 molecules. Error bars are s.e.m.

## Discussion

Our empirical finding demonstrates that BiP and ERdj3, the Sec61 tunnel-associated variants of DnaK and DnaJ, exhibit strong foldase activity that is practically relevant to their biological functions (Fig. 6). Since protein folding at the edge of any translocon pore act as a source of force generation (10, 19), these chaperones might influence the translocation of the substrates by promoting their folding at higher force range. We have also observed that both BiP-chaperone complex, and its individual component BiP and ERdj3 upshift the folding probability of protein L towards the higher force range (Fig. 2a and b), which further increases the force transmission through the Sec61 tunnel. As a feedback force-based mechanism, the translocation system opts for an optimum force transmission; which precisely tunes the intrinsic folding ability of substrate against the force, and thereby the generation of mechanical workdone. Our study revealed that BiP chaperone generate ∼54 zJ mechanical workdone by boosting the force output (or half-point force) from 6.4 to 7.7 pN, which plausibly aids in rescuing the arrested polypeptide within the Sec61 pore, and consequently their faster folding. Despite their strong foldase activity under mechanical constraints, both BiP and ERdj3 act as holdase in the near zero force, suggesting that mechanical behaviour of these chaperones is completely different from their foldase function at high force. We have observed that BiP chaperone complex also generate ∼1.7 times more mechanical energy than in its absence, indicating their active engagement in the force-mediated translocation process. Similarly, ERdj3 also delivers higher mechanical work output during protein L folding than its cytosolic variant DnaJ.

This mechanical protein folding by BiP chaperone complex not only triggers the accelerated folding but also prevent the backsliding of the polypeptide during the translocation through Sec61 translocon pore. To simply understand the physical basis of this co-translocational protein-folding, the tunnel-accommodated protein L can be assumed as a stretched polymer which is under effective force owing to the tunnel-imposed geometric constraints (37). In our study, we have found this effective force to be within 7-12 pN, which is physiologically relevant during translocation and also in agreement with the reported entropic pulling force (38). Protein folding at the tunnel-opening induces a shortening of the confined polypeptide, which facilitates the force-transmission through the tunnel and thus induces a molecular strain to hinder the protein folding. Considering the protein folding as a physical phenomenon, this strain will be stored as elastic potential energy within polypeptide; which must be released by it in a form of a mechanical workdone to attain a stable folded state. To model this, we have calculated the change in chaperone strain energy (SE) during their interaction with protein L, described as Eq. 3 (39, 40):

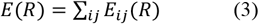

Where, *E* is potential energy of a configuration *R* which can be intrinsic deformation of the protein structure with *i* and *j* residue pair in the configuration. Now, if we consider the substrate-bound chaperone as a single system, the mechanical folding or unfolding of the substrate will bring about a consequent topological change to the chaperone causing its local deformation as well. To quantify the effect of this topological change on chaperone deformability, we introduced a parameter strain energy difference (ΔSE) as Eq. 4:

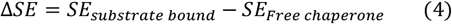

Here we observed DnaK and DnaJ exhibit higher deformability (or strain energy difference, Δ*SE*) with unfolded protein L and lower Δ*SE* while forming complex with folded state (Fig. 5d) unlike their respective homologs BiP and Erdj3, which exhibit higher deformability (or strain energy difference, Δ*SE*) with folded protein L and lower Δ*SE* with unfolded state (Fig. 5c). To relate this result with refolding ability of chaperones under mechanical influence, we introduced a second parameter called ΔΔ*SE*, calculated as Eq. 5:

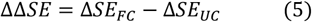

Where, Δ*SE*_*FC*_ and Δ*SE*_*UC*_ are strain energy difference while folded and unfolded substrates are bound to the chaperone, respectively. Since the intrinsic strain and thereby the substrate binding effect of chaperones vary, and can be compared only between structural homologs, we used normalized ΔΔ*SE* for comparative analysis of the chaperones (Eq. 6):

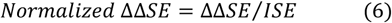

Interestingly, the tunnel-associated homologs BiP and ERdj3 exhibit a positive normalized ΔΔ*SE* (Fig. 5a), which consequently generates higher mechanical workdone than that of DnaK and DnaJ with negative normalized ΔΔ*SE* (Fig. 5b), during the protein L folding under force. This signifies that chaperones having negative normalized ΔΔ*SE* parameter that will retard the refolding of the stretched polypeptide, and thereby maintain at its unfolded state, whereas chaperones with positive normalized ΔΔ*SE* will aid refolding. Consensually, if ISE of the bound chaperone itself is higher, then the deformation caused by stretched substrate will promote more flexibility to the chaperone. This flexibility will induce freedom of movement in the chaperone leading to entropic unidirectional pulling of the substrate (38). Moreover, the transfer of substrate strain to the bound BiP/Erdj3 will help the substrate to fold faster by elastic means. By contrast, chaperones with lower intrinsic deformability cannot withstand the potential energy transferred by the substrate and therefore will lose it immediately to gain stability, leaving the substrate in its unfolded state. This model is strengthened by the fact that at no (or very low) force, BiP/Erdj3 and DnaK/DnaJ behave similarly (holdase), as the former will not get any freedom of movement and hence no entropic-pulling associated protein folding will occur. Since chaperone activity depends on substrate topology and size, we speculate that our result is specific for protein L and can possibly vary in the case of other substrates. Therefore, this increased folding probability and strain energy data indicate that BiP and ERdj3 shift the protein L folding dynamics towards the folded state through increasing their own internal strain energy, escalating the generation of mechanical energy which helps in active folding under force. Overall, this provides an in-depth understanding of mechanical chaperone activity through the internal strain energy that could potentially explains the unprecedented physical basis of chaperone-mediated protein folding under force.

**Figure 5:**
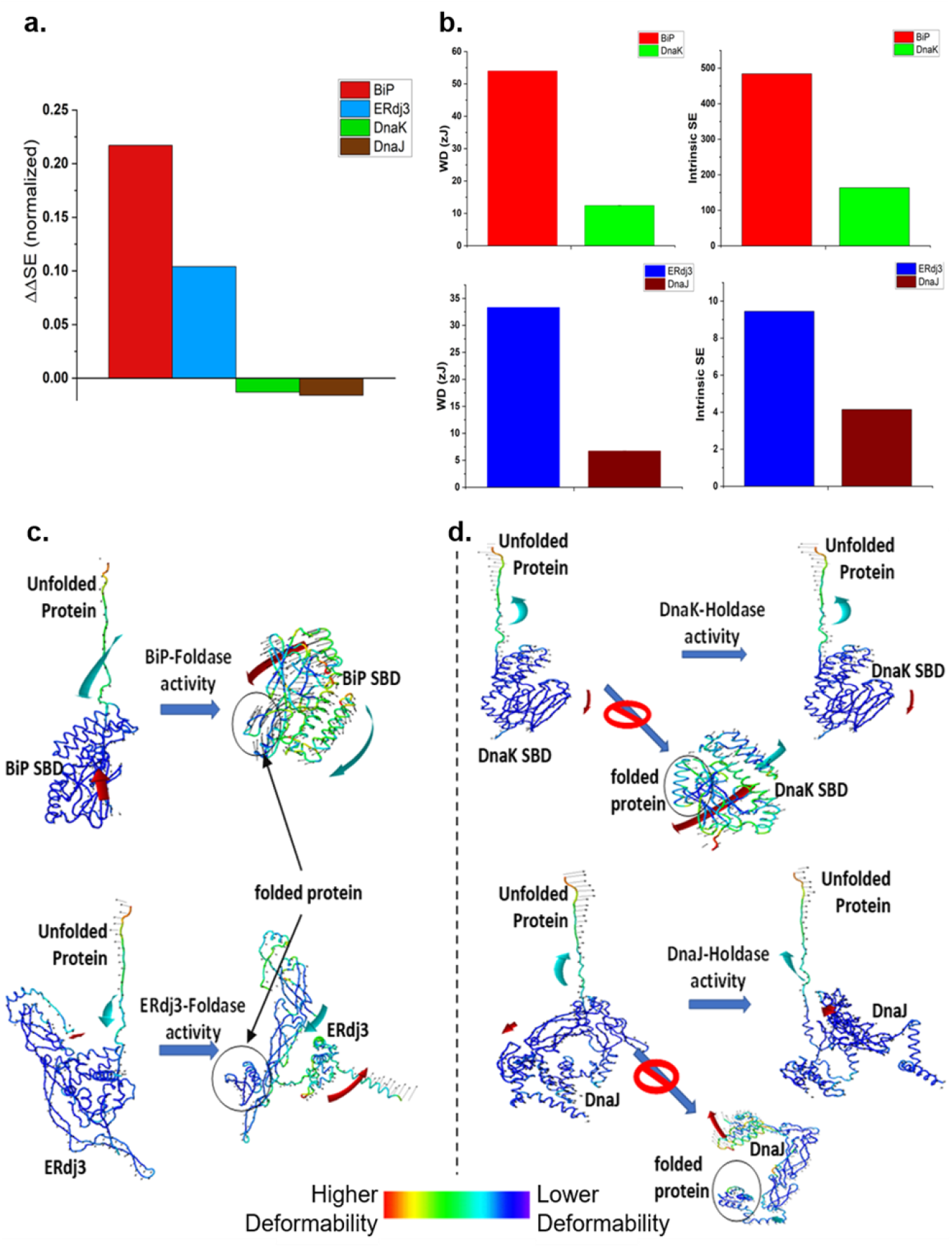
Intrinsic strain energy of chaperones regulates their mechanical activity during their interaction with substrate: **(a)** The strain energy difference (ΔSE) is obtained from intrinsic strain energy (ISE) of only chaperone and chaperon-substrate complex (ΔSE= ISE_C_ – ISE_SC_). Since, ΔSE value of foldase is higher with the folded state than that with unfolded; the difference in ΔSE value (or ΔΔSE= ΔSE_FC_ - ΔSE_UC_) will be positive for mechanical foldase chaperone. However, holdase chaperone has been observed to possess negative ΔΔSE. ISE of each chaperone are estimated from their SBD structures deposited in WEBnm@: BiP_SBD (PDB:5e85), DnaK_SBD (1dkz), DnaJ (AF-P08622-F1), and ERdj3 (AF-Q9UBS4-F1). **(b)** ISE of BiP is almost ∼4.5 times higher if compared to their holdases variants DnaK; which further could be correlated with their increased mechanical activity that BiP generate more workdone than DnaK at 7 pN (above). Similarly, we observed that ERdj3 has higher ISE than its holdase homolog DnaJ, which reflects into their ability to deliver higher mechanical work done. **(c and d)** BiP and ERdj3 bind to the unfolded protein L, in their lowest deformed state similar to their respective homolog DnaK and DnaJ (labelled as blue in all the chaperone structures). However, as the foldases contain higher ISE, they attain a higher deformability (labelled as green color) while transferring the potential energy from the unfolded protein L to them during the protein L folding. In contrast, DnaK and DnaJ have lower ISE and thus, cannot attain higher deformability during the folding of unfolded protein L. Consequently, these holdase chaperones stabilize the protein L at their unfolded conformation. Complexes are represented as heat maps according to region-specific deformability. Colors are indicative of their deformation energy as shown in figure. Small black arrows indicate the extent of deformation according to the specific direction and the large arrows show global changes. ISE_C_ is ISE of only chaperone; ISE_SC_ is ISE of chaperone-substrate complex. ΔSE_FC_ and ΔSE_UC_ are strain energy difference while folded and unfolded substrates are bound to the chaperone, respectively.

**Figure 6:**
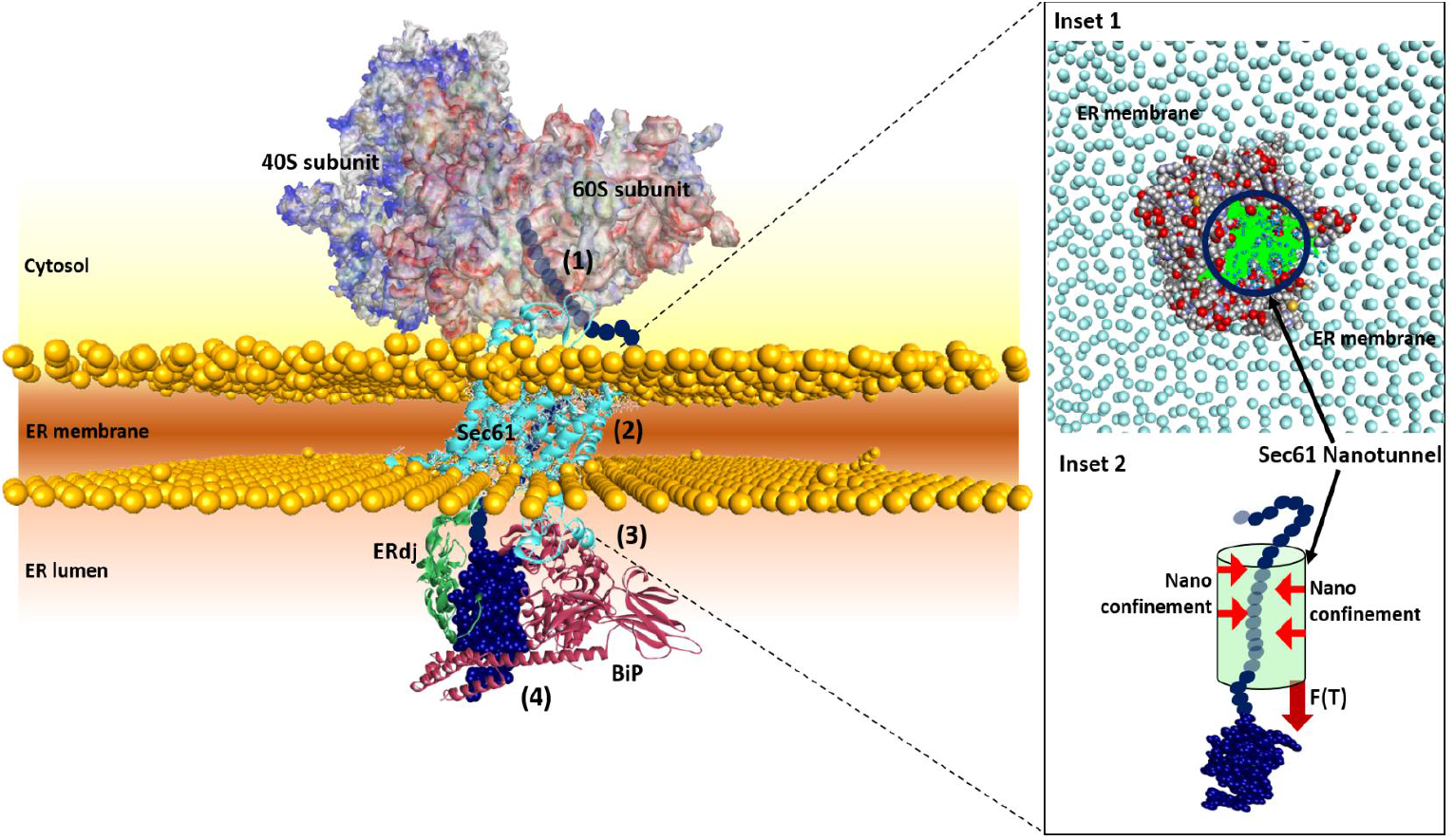
Model of BiP and ERdj3 as the mechanical foldase: During co-translational translocation of a nascent polypeptide chain through Sec61 tunnel of endoplasmic reticulum (ER), the chains are unable to fold itself within the tunnel due to their confined geometry (red arrows in inset 2) and repulsive electrostatic force of chain residues within it. However, upon translocating a significant fraction of it in the luminal face, the polypeptide at the edge of this tunnel pulls the unfolded peptide within the tunnel, transmitting the mechanical tension throughout the unfolded polypeptide. This folding of the polypeptide chain against the force results in the mechanical workdone. Sec61 tunnel-associated chaperones BiP (brown) and ERdj3 (green) bind to the emerging polypeptide at luminal face and pull the polypeptide to accelerate their folding against the force and thus, prevent their backsliding. Owing to the higher deformability of BiP and ERdj3 chaperones, they stabilize the folded state of the substrate and deliver higher mechanical workdone to hasten the protein folding during co-translational translocation.

## Materials and methods

### Protein L B1 domain octameric substrate expression and purification

The protein L octamer construct was subcloned using *Bam*HI, *Bgl*II and *Kpn*I restriction sites in pFN18A vector, as described previously (9). The construct is transformed into BL21(DE3 Invitrogen) *E. coli* and then cultured in Luria broth, supplemented with 50 μg/ml carbenicillin for the selection. The cultures were induced with 1 mM IPTG (Isopropyl β-D-thiogalactopyranoside) for overnight at 25ºC upon the O.D._600_ increases to 0.6-0.8. Then the cells were centrifuged and resuspended in 50 mM sodium (Na) phosphate buffer, supplemented with 10% glycerol and 300 mM NaCl (pH 7.4). Subsequently, Phenylmethylsulfonyl (PMSF) as protease inhibitor was added, followed by lysozyme for membrane lysis. This solution was incubated on ice for 20 mins and later triton-X100, DNase, RNase and MgCl_2_ were added and incubated gently on a rocking platform at 4 °C for 10 min. Cells were homogenized using cell disrupter at 19 psi; and finally, lysate was collected after centrifugation. The proteins were purified from the lysate through Ni^2+^-NTA affinity chromatography, followed by in vitro biotinylation and size exclusion chromatography (SEC).

### Expression and purification of chaperones

Human BiP (HSPA5; P11021) and human ERdj3 (DNAJB11; Q9UBS4) were cloned into pET30a+ vector with XhoI and NdeI. BiP and ERdj3 chaperones are transformed into BL21 (DE3) and RIL *E. coli* cells, followed by induction with 1 mM IPTG at 37º for 6 h and overnight at 25ºC, respectively. Both the proteins are purified by Ni-NTA affinity chromatography, followed by SEC. Similarly, after transforming into BL21 (DE3) cells, DnaK and DnaJ were grown in LB medium with respective antibiotics, centrifuged into pellet and resuspended in Na-phosphate buffer. Thereafter, the chaperones are purified according to the protocol described previously (17, 41).

### Single-molecule magnetic tweezers experiment

Magnetic tweezers experiments were carried out in a fluid chamber where two different size of coverslips were sandwiched using parafilm strips as the spacer. The glass coverslips were cleaned exactly as described previously (41). In short, the glass slides were washed with 1.5% Hellmanex III solution and washed with double-distilled water. The slides were then dipped in a solution of methanol and hydrochloric acid, followed by soaking into sulfuric acid and then washing. The slides were then treated in boiling water and then dried. For activating the glass surface 1% (3-Aminopropyl) trimethoxysilane was dissolved in ethanol and was silanized for 15 minutes. Then the glasses were washed with ethanol to remove nonspecific silane and baked at 65°C for ∼ 2 hour. Similarly, top coverslips were washed with 1.5% Hellmanex III solution for 15 minutes, followed by washing with ethanol and then drying it in the oven for 10 minutes. The chamber was then sandwiched using slide and coverslip, which was then flowed with glutaraldehyde (Sigma Aldrich, G7776) in PBS buffer. After an hour, the chamber was washed with non-magnetic beads (Spherotech, AP-25-10) followed by Halo-Tag amine (O_4_) ligand (Promega, P6741). The chambers were then treated with a blocking buffer (20 mM Tris-HCl of pH 7.4, 150 mM NaCl, 2 mM MgCl_2_, 1% BSA, and 0.03% NaN_3_) to avoid non-specific interaction and kept in room temperature for 5 hours. These reactive fluid chambers were used for further experimentations by mounting them on a stage, controlled by a nanofocusing piezo actuator (P-725; Physik Instrumente), of an inverted microscope with oil immersion objective (9, 16, 41). Above the fluid chamber permanent neodymium magnets are attached to a voice coil actuator which regulate the distance of the magnets finally controlling the applied force. Biotinylated protein L substrates were then allowed to react with the halo tag ligand which allowed them to covalently tether with the bottom glass. Streptavidin coated paramagnetic Dynabeads (M-270) were then passed through the fluid chamber to react with the biotinylated substrate proteins (9). All the experiments were carried out with 5 nM of protein L in PBS buffer (pH 7.4). Additionally, all the BiP, DnaK, and BiP complex experiments were performed in the presence of 10 mM MgCl_2_ and 5 mM ATP (replenished every 40 mins). For the experiments with BiP chaperone complex, its constituent chaperone (BiP, ERdj3, and Sil1) concentrations were chosen in accordance to the FP obtained at 7 pN.

### Boltzmann plot for the concentration-dependent change of folding probability at 7 pN

The concentration-dependent FP data points at 7 pN has been fitted using Boltzmann equation (Eq. 5).

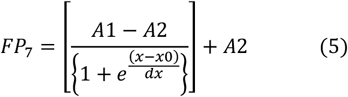

here, *FP*_*7*_ is *FP* at 7 pN of force at different concentrations, *A1*= initial FP, *A2*= final FP, *x0*= concentration at half the effect, and *dx*= time constant

### Calculation of %folded data obtained in near-zero force

The %folded data both in the presence and absence of chaperones were fitted as a function of quench time in near-zero force (0.3 pN) with exponential growth function:

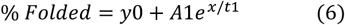

### Calculation of mechanical work done

The folding work at a particular force can be calculated by multiplying the force with step size. Then the expected value of mechanical workdone will be equal to the folding work multiplied by folding probability at a certain force.

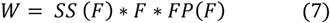

### Computational analysis of the mechanical properties of chaperone

To determine why BiP and ERdj3 behave as mechanical foldases while their cytoplasmic homologs do not, we analyzed the mechanical properties of all these chaperones. Firstly, we considered the structures of the substrate binding domain of BiP (5e85) and DnaK (1DKZ) (Supplementary Fig. 19). The folded conformation of protein L (1HZ6) was chosen and unfolded conformation of protein L substrate was prepared using steered molecular dynamics simulation in NAMD, as described previously (41). Only AlphaFold structures of ERdj3 and DnaJ were chosen due to unavailability of their crystallographic structures.

The intrinsic deformability was calculated for each of these chaperones using WEBnm@ web-server, which calculates deformation energy of different normal modes conformations defined by their eigenvalues. The mean of deformation energies of the closest non-trivial normal modes was chosen for the chaperones in each case. Such as, for DnaJ the value of deformation energy of mode 7 was 4.1 whereas mode 8 value was 8.01, so we proceeded further for DnaJ with mode 7 data only. However, its homologue ERdj3 showed DE values of 7.3, 9.26, 9.7 and 11.59 in mode 7 to 10 before jumping to 23.35 in mode 11. So, for ERdj3 the mean DE of mode 7 to 10 was considered as the DE or the intrinsic strain energy (SE). Similar way was used to identify the intrinsic SE for DnaK_SBD and BiP_SBD. These chaperones were then docked with folded and unfolded protein L substrate using HADDOCK 2.4 (Supplementary Fig. 22 and 23) (42) after obtaining the maximum confidence score of the interacting molecules, using HDOCK (43). The results obtained from HADDOCK 2.4 were represented as van der Waals energy and electrostatic energy as provided (Supplementary Fig. 20 and 21).

### Analysis

All the data acquisition and analysis were performed with Igor Pro 8.0 software (Wavemetrics) and Origin software. Simulation data are analyzed with Biovia Discovery studio visualizer.

## Author contribution

S.H. and S.B. designed the project. S.B., D.C., and S.H performed the experiments. S.B., S.H.,

D.C and S.C. wrote the manuscript.

## Acknowledgments

We thank Ashoka University, Trivedi School of Biosciences and Mphasis foundation for support and funding. S.H. thanks DBT Ramalingaswami Fellowship and DST SERB Core Research Grant for funding.

## Supplementary Information

**Supplementary Fig 1:**
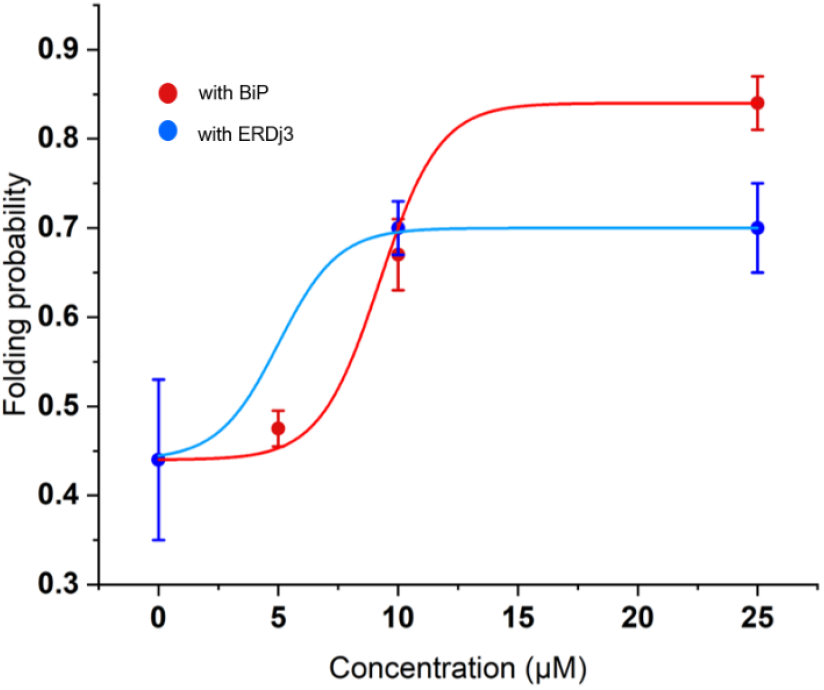
Concentration-dependent change in protein L folding probability: Folding probability of protein L has been measured with different concentrations of BiP and ERdj3, at 7 pN force. Folding probability of protein L changes significantly while increasing the ERdj3 concentration upto 10 μM; however, becomes stable after further increasing its concentration to 25 μM. Similarly, with BiP, the folding probability increases upon increasing the concentration to 10 μM but remains stable at its 25 μM concentration. Each data point represents n≥4 molecules and were calculated using > 250 s for every concentration. Error bars are standard error of mean (s.e.m).

**Supplementary Fig 2:**
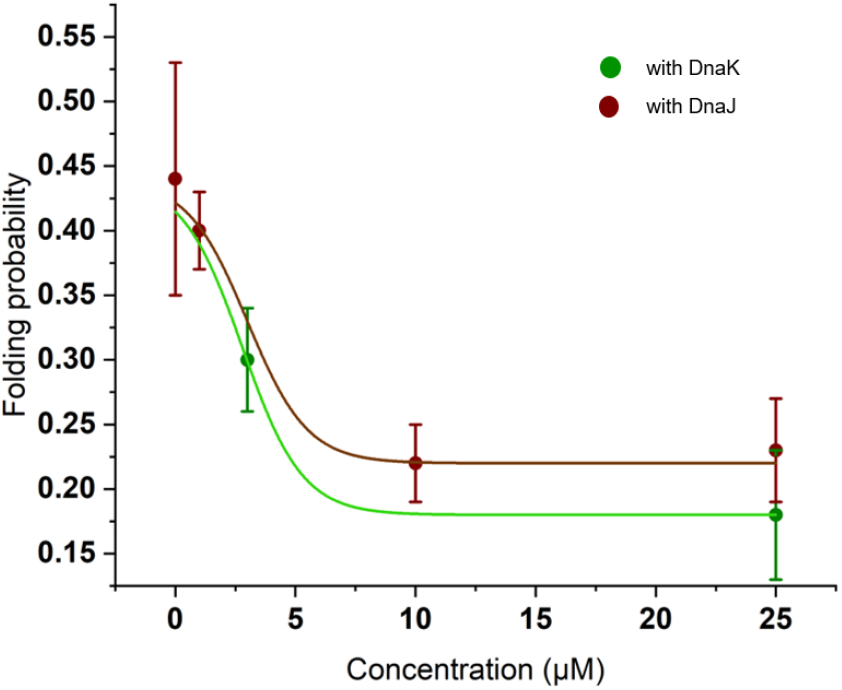
Concentration-dependent change in protein L folding probability in presence of DnaK and DnaJ: Folding probability of protein L has been measured with different concentrations of DnaK and DnaJ at 7pN force. Both these chaperones were observed to negatively affect the FP of protein L after gradual increase of the concentrations till the FP saturates. Folding probability of protein L changes significantly while increasing the DnaJ concentration upto 10μM; however, becomes stable after further increasing its concentration to 25μM. Similarly, with DnaK, the folding probability decreases upon increasing the concentration. Each data point represents n≥4. Error bars are s.e.m.

**Supplementary Fig 3:**
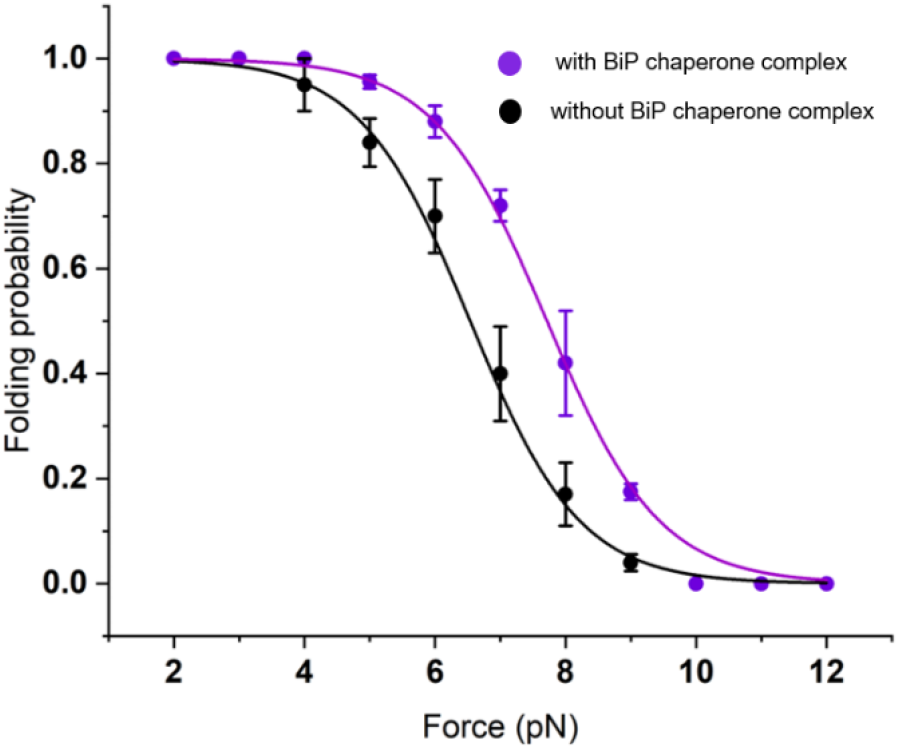
Folding probability in presence of the BiP chaperone complex: BiP chaperone complex (containing 25 μM BiP, 10 μM ERdj3, and 10 μM Sil1; in addition to 5 mM ATP and 10 mM MgCl_2_) was used to check its effect on the folding probability of protein L. We have observed that BiP chaperone complex increases the folding probability of protein L by upshifting the half-point force from 6.4 pN to 7.7 pN. Each data point represents n≥5 molecules and were observed for >250 s. Error bars are s.e.m.

**Supplementary Fig 4:**
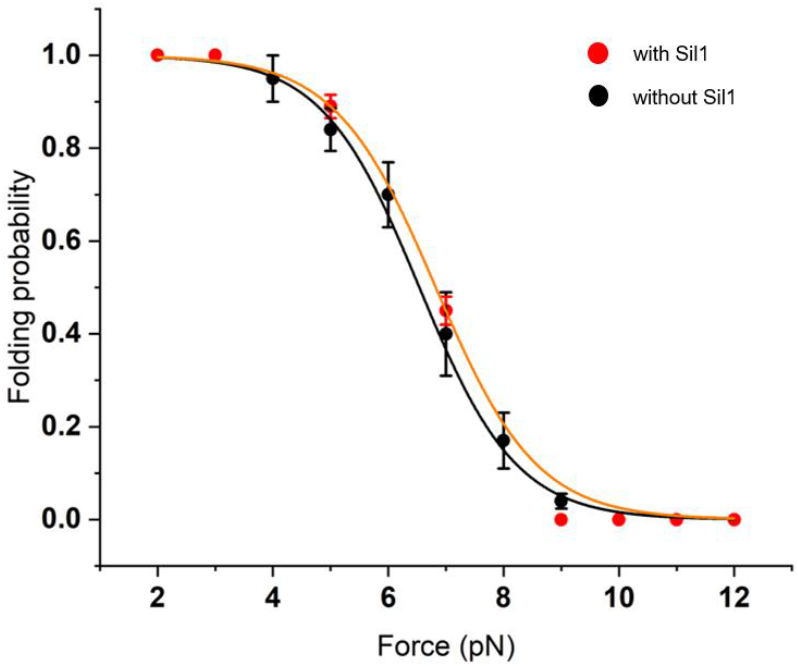
Folding probability of protein L with Sil1: Folding probability of protein L at different forces has been plotted to understand the effect of Sil1. We observed that the folding probability of protein L with 10 μM Sil1/BAP (red circles) is overlapped with that in their absence (control, black circles) with a similar half point force of 6.7 pN. Each data point represents n≥4 molecules. Each datapoints of FP were calculated using >250 s. Error bars are s.e.m.

**Supplementary Fig 5:**
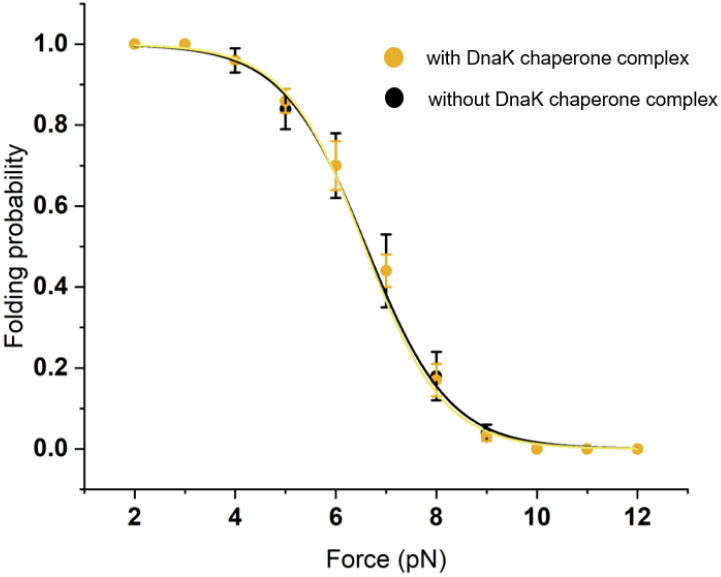
Folding probability of protein L with DnaK chaperone cycle: Folding probability of protein L at different forces has been plotted to understand the effect of DnaK chaperone complex. We observed that the FP of protein L in presence of 25μM DnaK, 10μM DnaJ and 10μM GrpE (yellow circles) overlaps with that in their absence (control, black circles) with a similar half point force of 6.6 pN. The experiments were performed in the presence of to 5 mM ATP and 10 mM MgCl_2_. Each data point represents n≥4. Error bars are s.e.m.

**Supplementary Fig 6:**
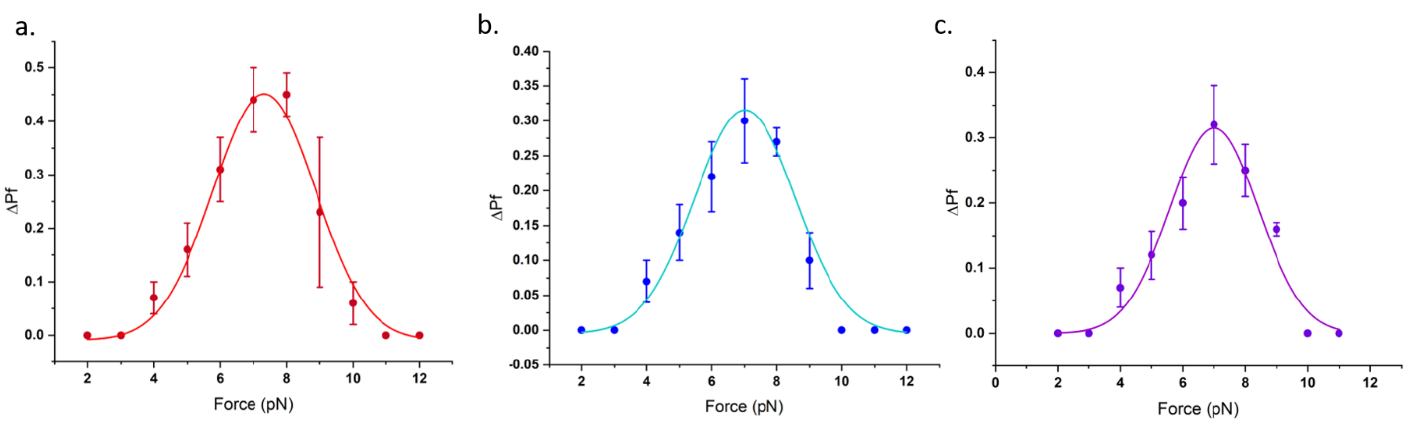
Relative increment in the folding probability of protein L in the presence of chaperones: Relative increment in the folding probability (defined as ΔPf) has been plotted as a function of force. **(a)** The ΔPf reaches to ∼45%. in the presence of BiP chaperone (red circles). Similarly, for **(b)** ERdj3 (blue), and **(c)** BiP chaperone complex (purple), the ΔPf reaches upto ∼30% increase at 7 pN of force. Each FP datapoints at every force were observed for >250 s. Each data point represents n≥4. Error bars are s.e.m.

**Supplementary Fig 7:**
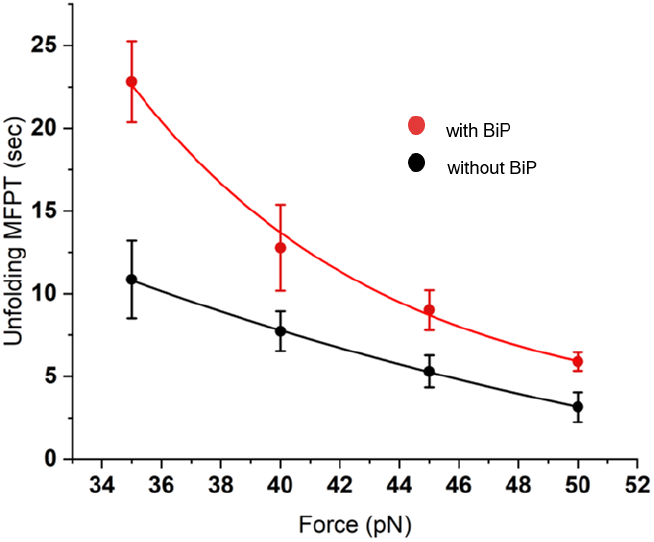
Unfolding MFPT in presence and absence of BiP: Unfolding MFPT has been plotted as a function of varying forces. At a particular force, the unfolding MFPT has been observed to increases in the presence of 25 μM BiP (red circles) than in its absence (black circles), indicating that BiP slows down the unfolding kinetics of protein L. Each data point represents n>4 molecules. Error bars are s.e.m.

**Supplementary Fig 8:**
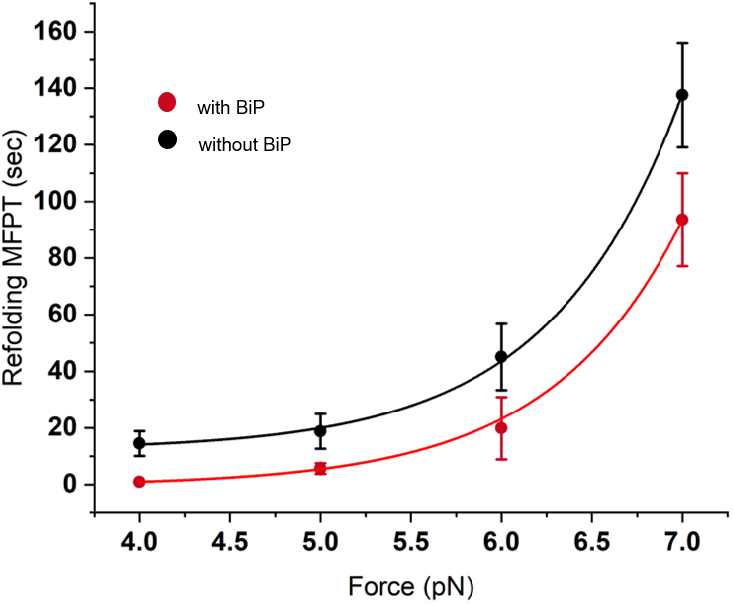
BiP accelerates the refolding kinetics: The refolding MFPT were plotted both in the absence (black circles) and presence (red circles) of BiP. We have observed that BiP accelerates the refolding kinetics by decreasing the total refolding time. For example, at 7 pN force, the polyprotein completely refolds in 137.65±18.1 sec without BiP; however, it decreases to 93.5±16.1 sec in the presence of BiP. Each data points represents n≥4 molecules and error bars are s.e.m.

**Supplementary Fig 9:**
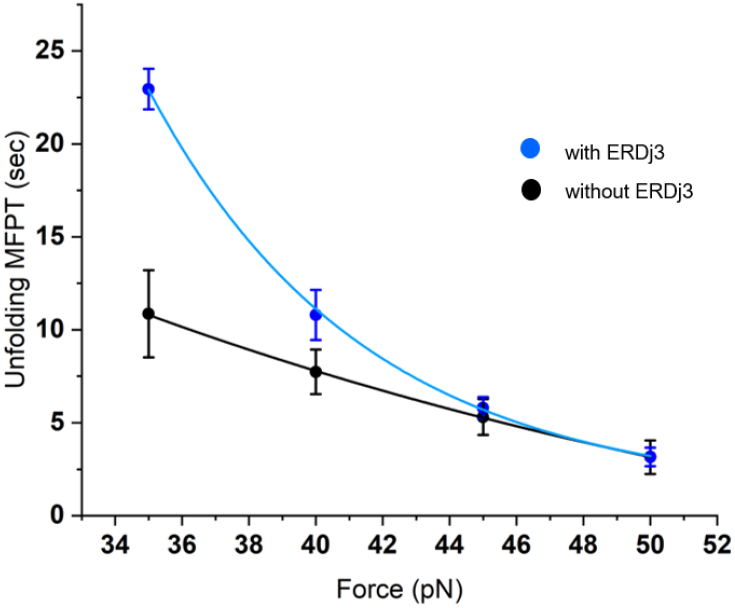
Unfolding MFPT in presence and absence of ERdj3: Unfolding MFPT has been plotted against the forces both in the absence (black circles) and presence of ERdj3 (blue circles). At a particular force, the unfolding MFPT has been observed to increases in the presence of 10 μM ERdj3 than in its absence, indicating that ERdj3 slows down the unfolding kinetics of protein L. Each data point represents n≥5 molecules. Error bars are s.e.m.

**Supplementary Fig 10:**
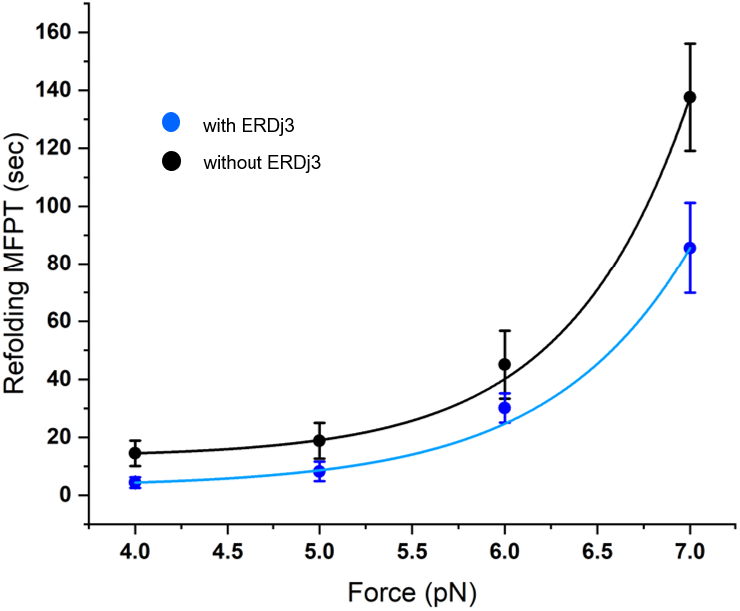
Refolding MFPT in the presence and absence of ERdj3: The refolding MFPT against the force has been fitted to exponential equation. We have observed that the refolding MFPT decreased from 137.65±18.1 sec without ERdj3 (control, black circles) to 85.56±15.6 sec in the presence of ERdj3 (blue circles). Each data point represents n>5 molecules and error bars are s.e.m.

**Supplementary Fig 11:**
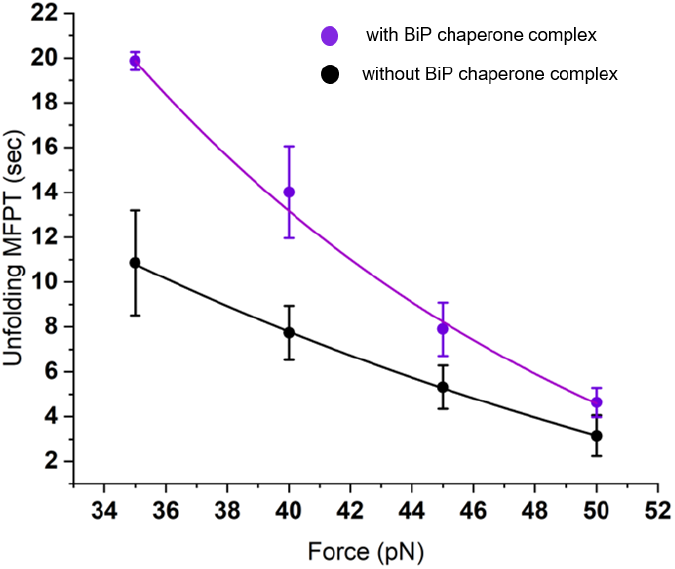
Unfolding MFPT with and without BiP chaperone complex (BiP, ERdj3 and Sil1): Unfolding MFPT has been plotted as a function of varying forces. At a particular force, the unfolding MFPT has been observed to increases in the presence of 25μM BiP, 10μM ERdj3 and 10μM Sil1 (violet circles) than in its absence (black circles), indicating that this chaperone complex slows down the unfolding kinetics of protein L. Experiments were done in presence of 5mM ATP and MgCl_2_. Error bars are standard error of mean (s.e.m). Each data point represents n>4.

**Supplementary Fig 12:**
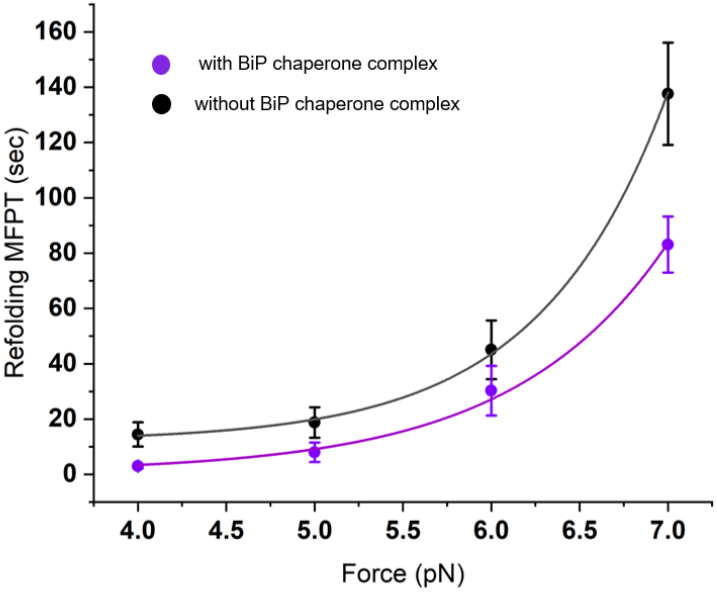
BiP chaperone complex accelerates the refolding kinetics: The refolding MFPT were plotted both in the absence (black circles) and presence (violet circles) of BiP chaperone complex. We have observed that BiP chaperone complex accelerates the refolding kinetics by decreasing the total refolding time. For example, at 7 pN force, the polyprotein completely refolds in 137.65±18.5 sec without BiP; however, it decreases to 83.1±10.4 sec in the presence of this chaperone complex. Each data points represents n≥4. Error bars are s.e.m.

**Supplementary Fig 13:**
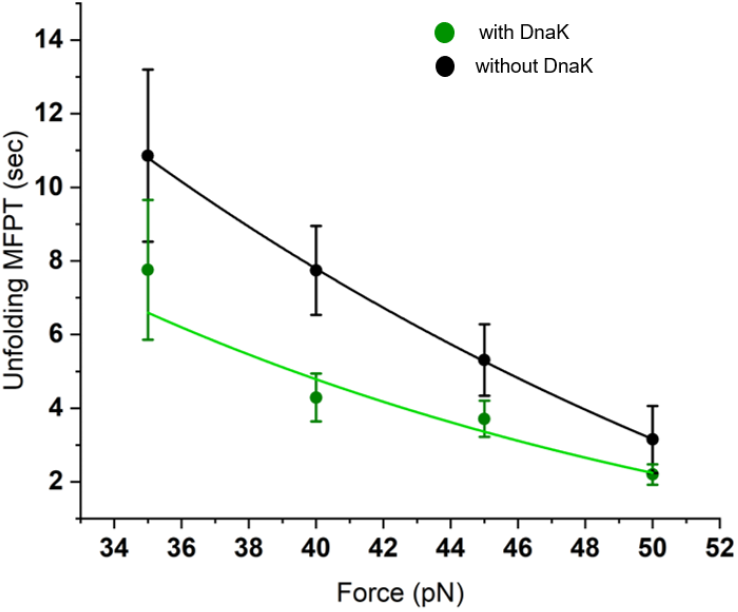
DnaK regulating unfolding MFPT of protein L: DnaK reduces the unfolding MFPT of protein L substrate as shown in this figure. In presence of 25μM DnaK (green circles) the unfolding of the polypeptide was occurring faster than when it is absent (black circles). Error bars are standard error of mean (s.e.m). Each data point represents n≥5.

**Supplementary Fig 14:**
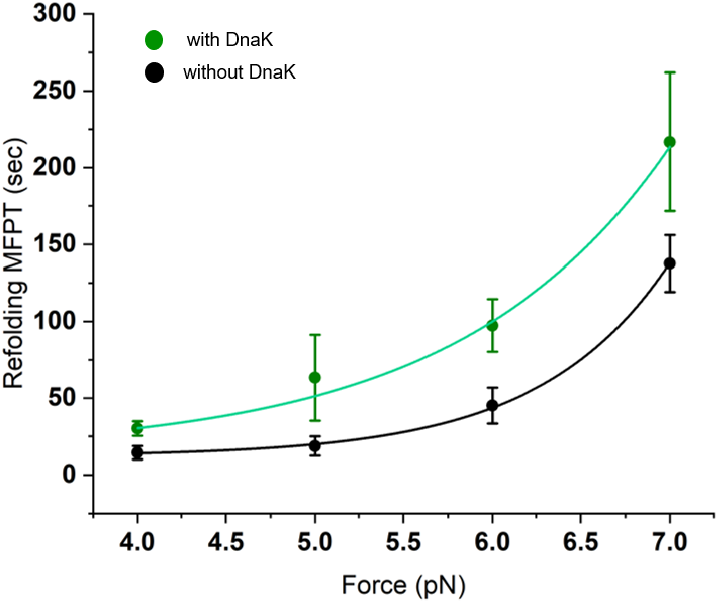
Refolding MFPT in the presence and absence of DnaK: Exponential fitting of the refolding MFPT against the force shows that refolding MFPT increased significantly in the presence of DnaK (green circles). Each data point represents n>5. Error bars are s.e.m.

**Supplementary Fig 15:**
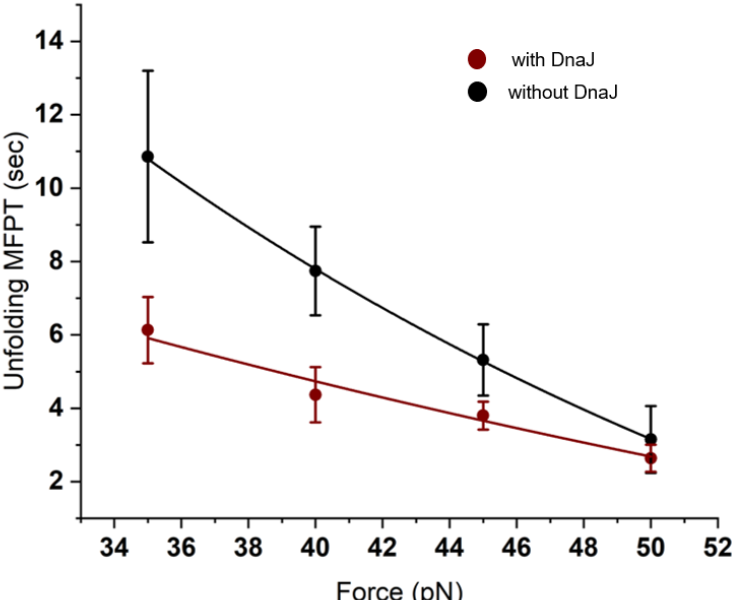
DnaJ decreases the unfolding time: By exponentially fitting the unfolding MFPT at different forces it was observed that DnaJ reduces the unfolding time of the substrate significantly. The data has been plotted against the forces both in the absence (black circles) and presence of 10μMDnaJ (brown circles). Error bars are standard error of mean (s.e.m). Each data point represents n≥5. Error bars are s.e.m.

**Supplementary Fig 16:**
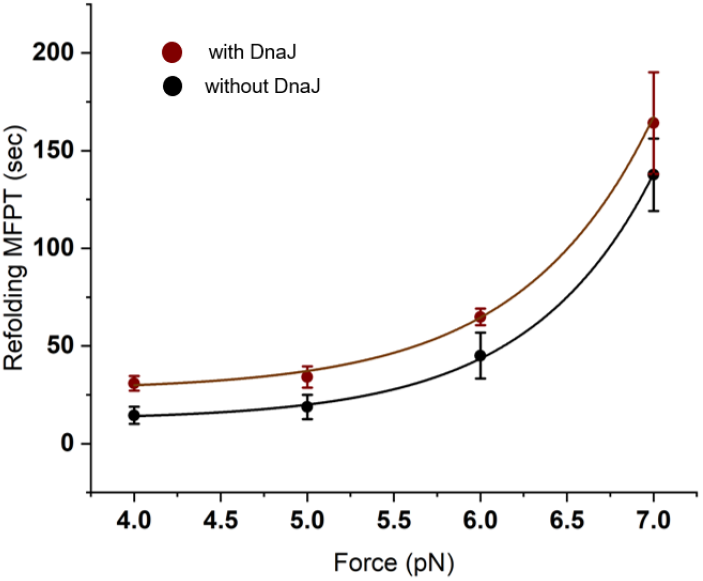
Refolding MFPT decelerates in presence of 10μM DnaJ: The refolding MFPT data is plotted against the force led to observe that it increased significantly in the presence of DnaJ (brown circles) and hence more time was required for proper refolding. Each data point represents n>5. Error bars are s.e.m.

**Supplementary Fig 17:**
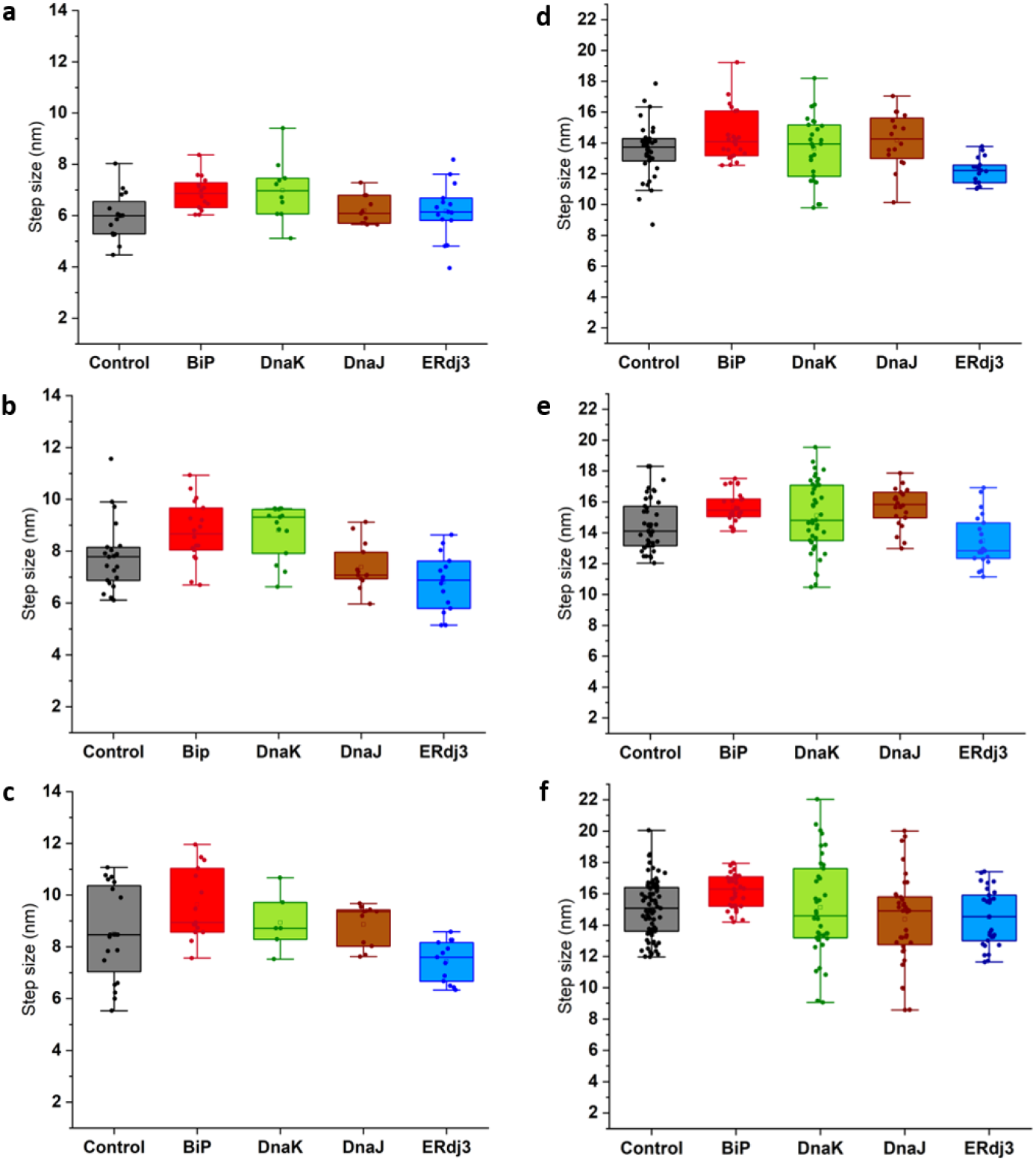
Comparative analysis of step size of protein L in presence of different chaperones at different forces: At each of the particular forces (a) 6pN, (b) 7pN, (c) 8pN, (d) 35pN, (e) 40pN, (f) 45pN; we analyzed the comparison in the step size of protein L in presence of different chaperone. No significant changes were observed in the step size of substrate protein at each force in presence of different chaperones.

**Supplementary Fig 18:**
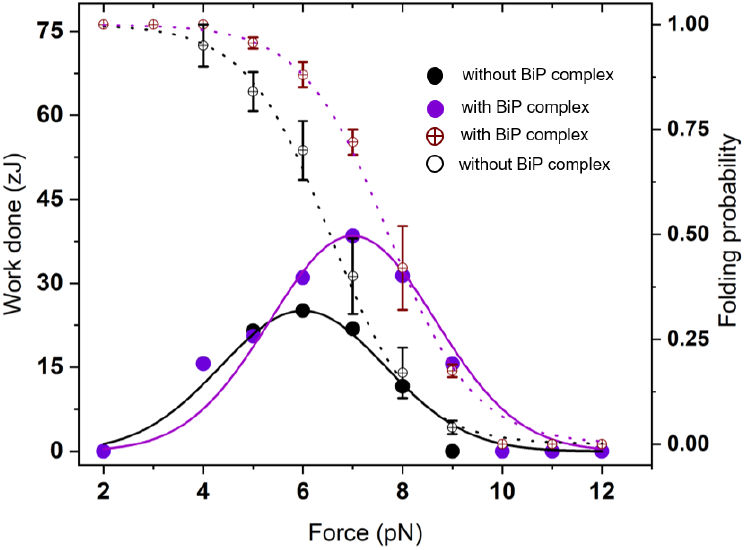
Mechanical workdone by the BiP chaperone complex: We measured the mechanical work done by the protein L in the presence of BiP chaperone complex and observed that mechanical work done of folding increases from 22 ±0.09 zJ in the control (black solid circles) to 38±0.013 zJ in the presence of BiP chaperone complex (violet solid circles) at 7 pN force.

**Supplementary Fig 19:**
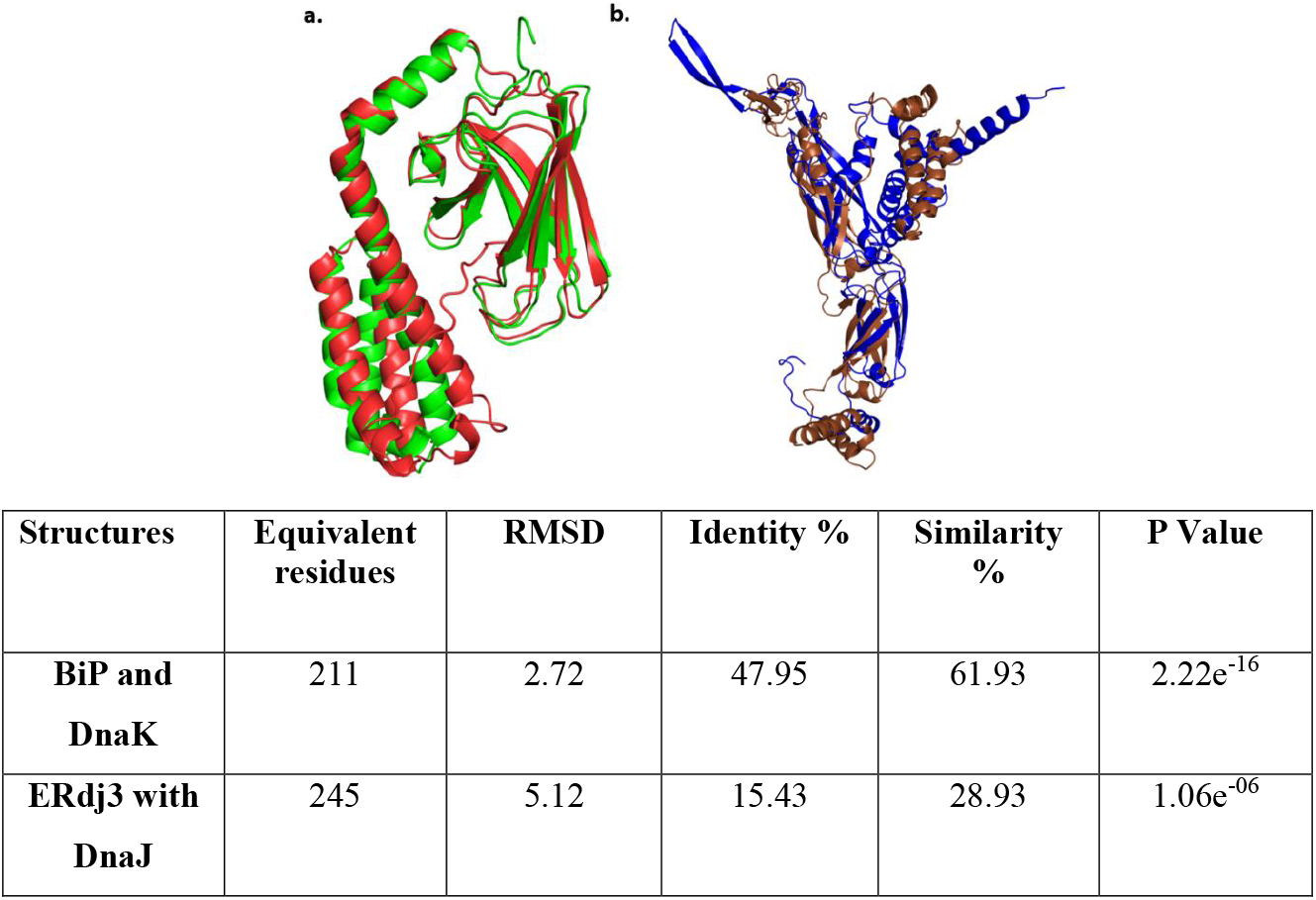
Structural alignment of substrate binding domain (SBD) of mechanical foldases with their cytoplasmic holdase variants: (a) SBD of BiP and DnaK: The structural alignment was done between BiP_SBD (red; PDB id: 5e85) and DnaK_SBD (green; PDB id: 1dkZ) using the webserver POSA **(Ref). (b) SBD of ERdj3 and DnaJ:** While aligning DnaJ (brown; AF-P08622-F1-model_v4) and ERdj3 (blue; AF-Q9UBS4-F1-model_v4) we proceeded with the Alpha fold structures as their X-ray crystallographic structures are not available. The structural alignment data observed with POSA has been provided in the table below.

**Supplementary Fig 20:**
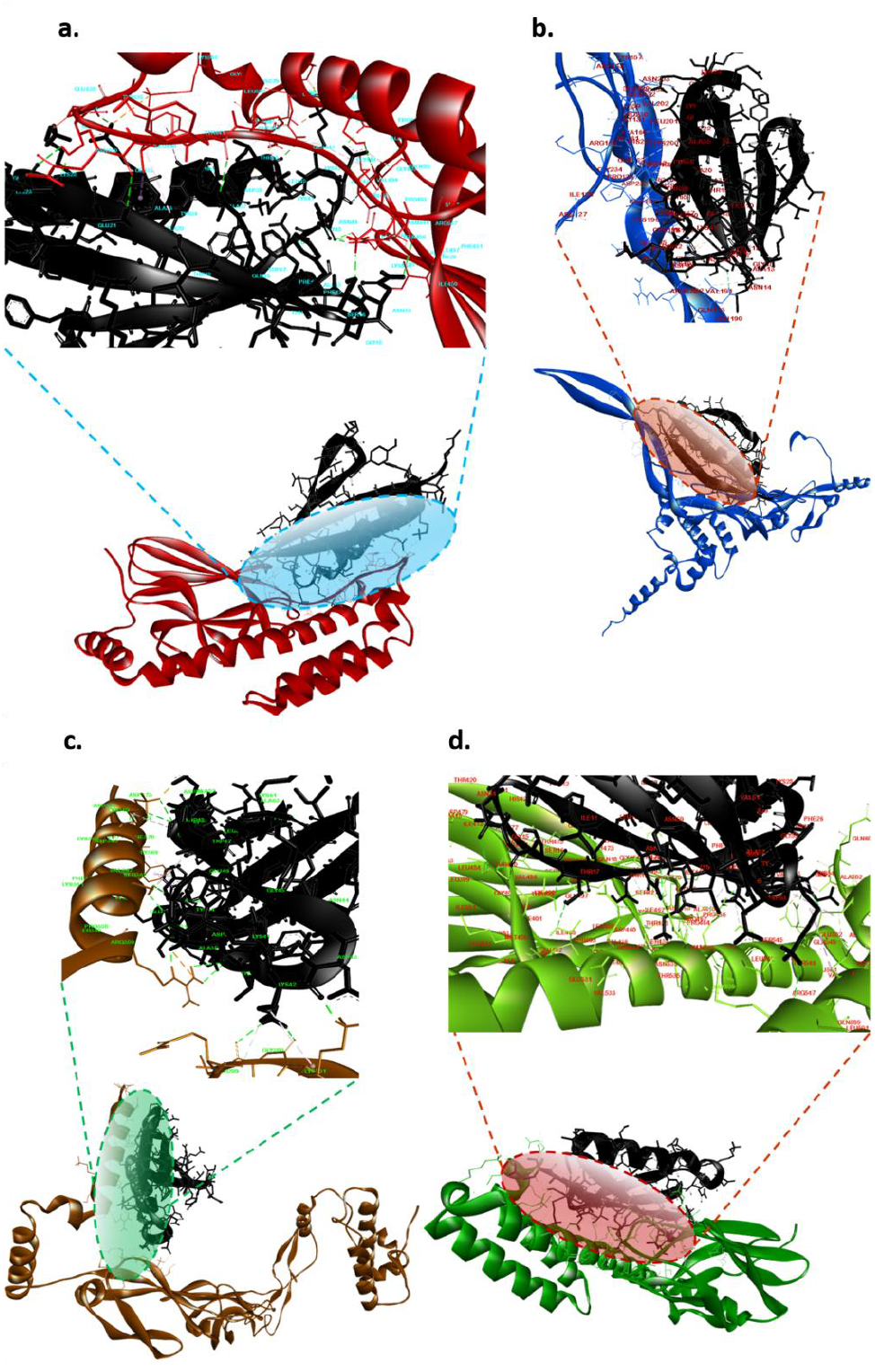
Docking analysis: **(a)** BiP_SBD (red), **(b)** ERdj3(blue), **(c)** DnaJ (brown), and **(d)** DnaK_SBD (green) with folded conformation of protein L B1 domain (black). The interacting amino acids are highlighted in the inset.

**Supplementary Fig 21:**
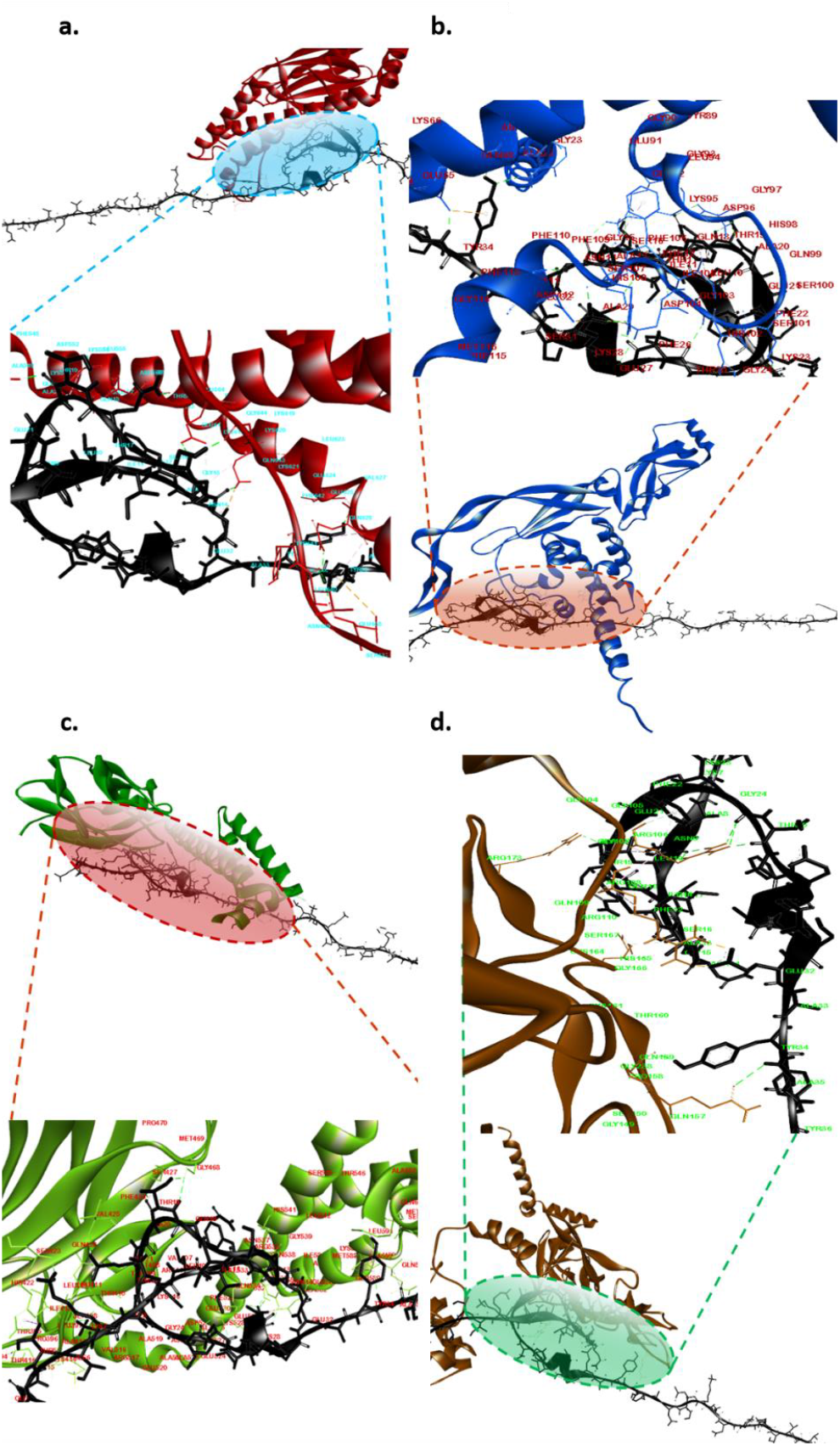
Docking analysis: **(a)** BiP_SBD (red), **(b)** ERdj3(blue), **(c)** DnaK_SBD (green), and **(d)** DnaJ (brown) with unfolded conformation of protein L B1 domain (black). The insets highlight the interacting amino acids of both the proteins.

**Supplementary Fig 22:**
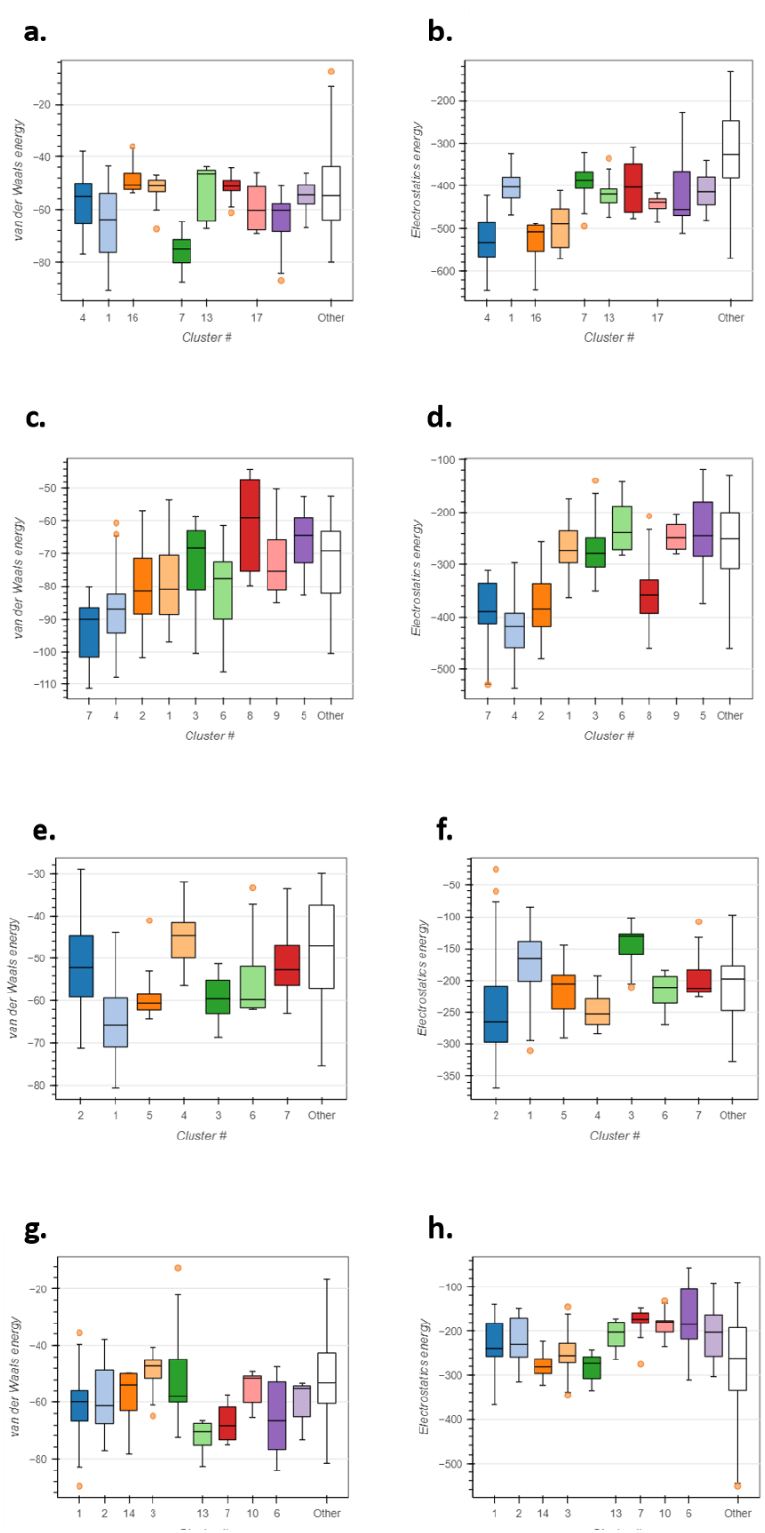
Binding energy of chaperones with folded substrate: For docking of mechanical foldases and holdases with folded protein L conformation using HADDOCK 2.4, we used water-refined models for all these molecules. Each cluster-generated complex structure representing van der Waals and electrostatic energy. The first cluster was selected for further analysis due to their lowest free energy and frequency. **(a, b)** BiP_SBD; **(c, d)** DnaK_SBD; **(e, f)** ERdj3 and **(g, h)** DnaJ docked with folded proteinL B1 domain.

**Supplementary Fig 23:**
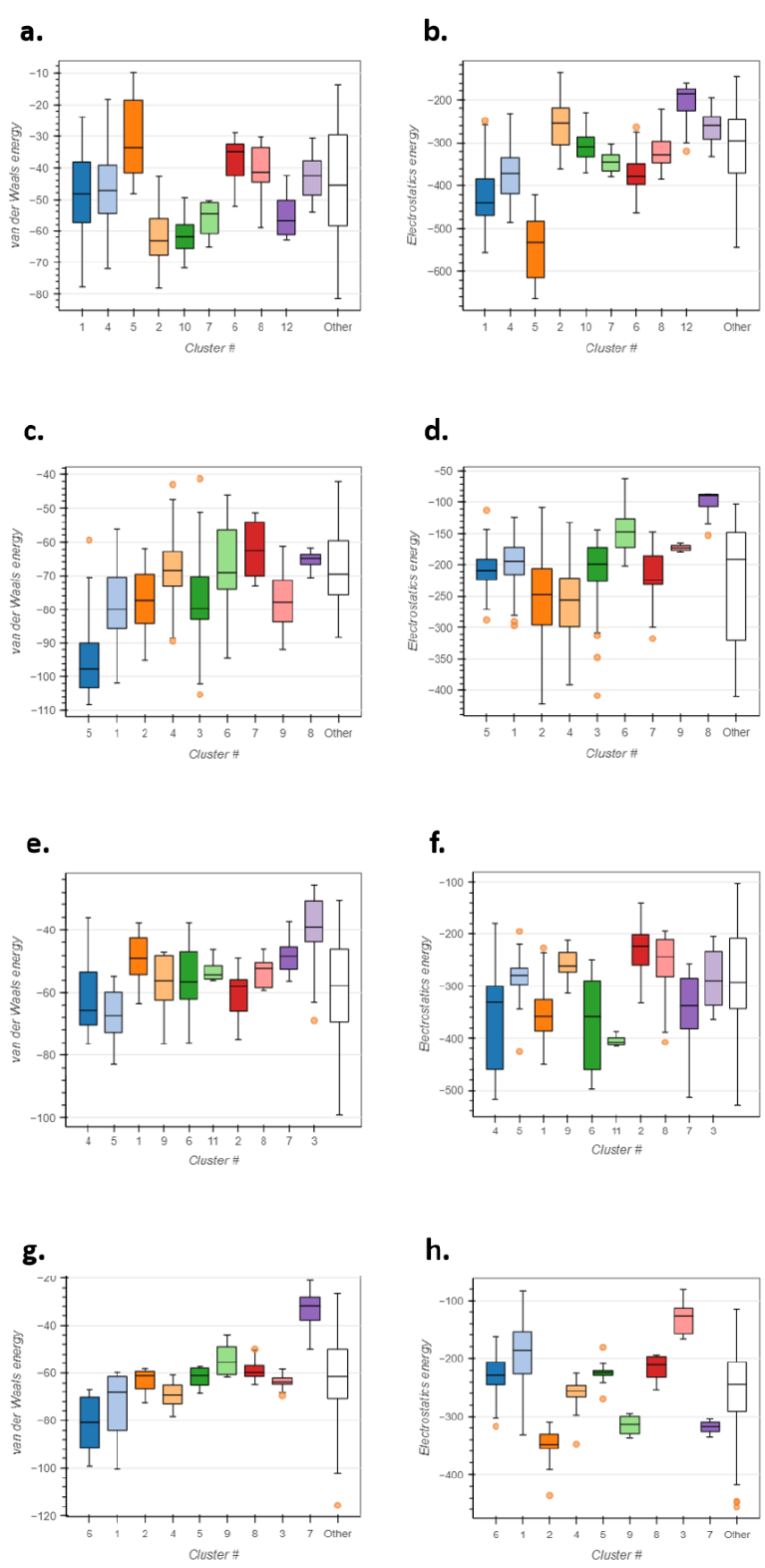
Binding energy of chaperones with unfolded substrate: Using HADDOCK 2.4, we docked both mechanical foldases and holdases with unfolded protein L conformation. The generated structures representing van der Waals and electrostatic energy according to the number of clusters generated during docking. The first cluster was selected for further analysis due to their lowest free energy and frequency. **(a, b)** BiP_SBD; **(c, d)** DnaK_SBD; **(e, f)** ERdj3 and **(g, h)** DnaJ docked with folded proteinL B1 domain.

